# A generic multivariate framework for the integration of microbiome longitudinal studies with other data types

**DOI:** 10.1101/585802

**Authors:** Antoine Bodein, Olivier Chapleur, Arnaud Droit, Kim-Anh Lê Cao

## Abstract

Simultaneous profiling of biospecimens using different technological platforms enables the study of many data types, encompassing microbial communities, omics and meta-omics as well as clinical or chemistry variables. Reduction in costs now enables longitudinal or time course studies on the same biological material or system. The overall aim of such studies is to investigate relationships between these longitudinal measures in a holistic manner to further decipher the link between molecular mechanisms and microbial community structures, or host-microbiota interactions. However, analytical frameworks enabling an integrated analysis between microbial communities and other types of biological, clinical or phenotypic data are still in their infancy. The challenges include few time points that may be unevenly spaced and unmatched between different data types, a small number of unique individual biospecimens and high individual variability. Those challenges are further exacerbated by the inherent characteristics of microbial communities-derived data (e.g. sparsity, compositional).

We propose a generic data-driven framework to integrate different types of longitudinal data measured on the same biological specimens with microbial communities data, and select key temporal features with strong associations within the same sample group. The framework ranges from filtering and modelling, to integration using smoothing splines and multivariate dimension reduction methods to address some of the analytical challenges of microbiome-derived data. We illustrate our framework on different types of multi-omics case studies in bioreactor experiments as well as human studies.

## 2 Introduction

Microbial communities are highly dynamic biological systems that cannot be fully investigated in snapshot studies. The decreasing cost of DNA sequencing has enabled longitudinal and time-course studies to record the temporal variation of microbial communities (Faust *et al.*, 2015; Knight *et al.*, 2012). These studies can inform us about the stability and dynamics of microbial communities in response to perturbations or different conditions of the host or their habitat. They can also capture the dynamics of microbial interactions (Ridenhour *et al.*, 2017; Bucci *et al.*, 2016) or associated changes of microbial features, such as taxonomies or genes, to a phenotypic group (Metwally *et al.*, 2018).

However, besides the inherent characteristics of microbiome data, including sparsity, compositionality (Aitchison, 1982; Gloor *et al.*, 2017), its multivariate nature and high variability (Lê Cao *et al.*, 2016b), longitudinal studies suffer from irregular sampling and subject drop-outs. Thus, appropriate modelling of the microbial profiles is required, for example by using splines modelling. Methods including loess (Shields-Cutler *et al.*, 2018), smoothing spline ANOVA (Paulson *et al.*, 2017), negative binomial smoothing splines (Metwally *et al.*, 2018) or gaussian cubic splines (Luo *et al.*, 2017) were proposed to model dynamics of microbial profiles across groups of samples or subjects. The aim of these approaches is to make statistical inferences about global changes of differential abundance across multiple phenotypes of interest, rather than at specific time points. These proposed methods are univariate, and as such, cannot infer ecological interactions (Morris *et al.*, 2016). Other types of methods aim to cluster microbial profiles to posit hypotheses about symbiotic relationships, interaction or competition. For example Baksi *et al.* (2018) used a Jenson Shannon Divergence metric to visually compare metagenomic time series.

Multivariate ordination methods can exploit the interaction between microorganisms, but need to be used with sparsity constraints, such as 𝓁_1_ regularization (Tibshirani, 1996), to reduce the number of variables and improve interpretability through variable selection. Several sparse methods were proposed and applied to microbiome studies, such as sparse linear discriminant analysis (Clemmensen *et al.*, 2011) and sparse Partial Least Squares Discriminant Analysis (sPLS-DA, Lê Cao *et al.* 2016a), but for a single time point. Therefore, further developments are needed to combine time-course modelling with multivariate approaches to start exploring microbial interactions and dynamics.

In addition, current statistical methods have mainly focused on a single microbiome dataset, rather than the combination of different layers of molecular information obtained with parallel multi-omics assays performed on the same biological samples. Data derived from each omics technique are typically studied in isolation, and disregard the correlation structure that may be present between the multiple data types. Hence, integrating these datasets enables us to adopt a holistic approach to elucidate patterns of taxonomic and functional changes in microbial communities across time. Some sparse multivariate methods have been proposed to integrate omics and microbiome datasets at a single time point and identify sets of features (multi-omics signatures) across multiple data types that are correlated with one another. For example, Gavin *et al.* (2018) used the DIABLO method (Singh *et al.*, 2019) to integrate 16S amplicon microbiome, proteomics and metaproteomics data in a type I diabetes study, Guidi *et al.* (2016) used sparse PLS (Lê Cao *et al.*, 2008) to integrate environmental and metagenomic data from the Tara Oceans expedition to understand carbon export in oligotrophic oceans, and Fukuyama *et al.* (2017) used sparse Canonical Correlation Analysis (Witten *et al.*, 2009) to integrate 16S and metagenomic data. However, methods or frameworks to integrate multiple longitudinal datasets including microbiome data remain incomplete. Zhou *et al.* (2008) used Principal Component Analysis (PCA) to summarize functional data, with the PC scores used for model fitting, prediction and inference. However, only pairwise relationships were investigated and for a single type of data. Other type of modelling (loess regression) was used by Ribicic *et al.* (2018) in combination with sparse PCA to explore the link between chemistry and microbial community data in the biodegradation of chemically dispersed oil, but their approach was not designed to seek for multi-omics signatures.

We propose a computational approach to integrate microbiome data with multi-omics datasets in longitudinal studies. Our framework, described in Figure 1 includes smoothing splines in a linear mixed model framework to model profiles across groups of samples, and builds on the ability of sparse multivariate ordination methods to identify sets of variables highly associated across the data types, and across time. Our framework encompasses data pre-processing, modelling, data clustering and integration. It is highly flexible in handling one or several longitudinal studies with a small number of time points, to identify groups of taxa with similar behaviour over time, and posit novel hypotheses about symbiotic relationships, interactions or competitions in a given condition or environment, as we illustrate in two case studies.

**Figure 1:**
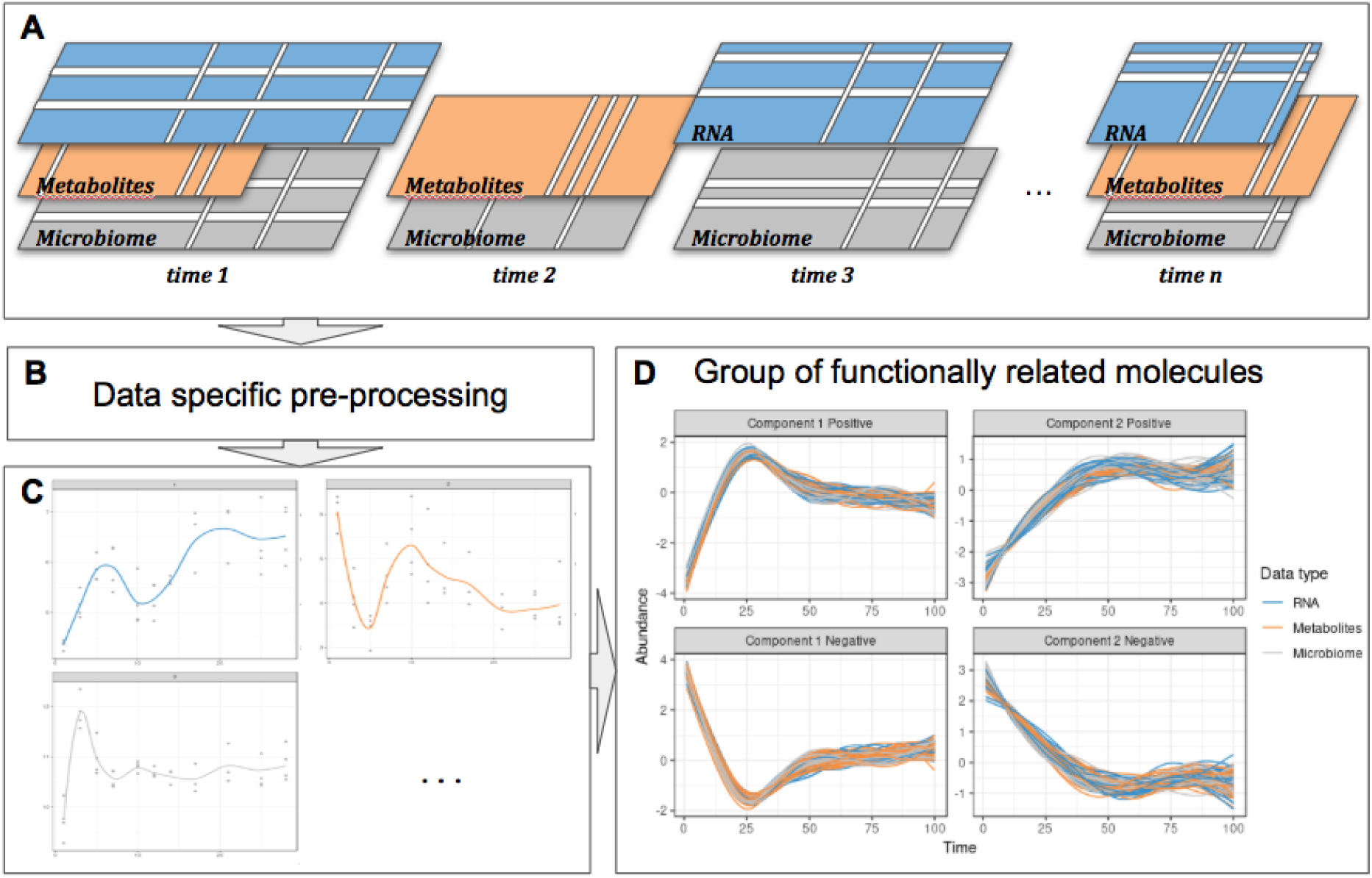
Overview of the proposed approach. **A**: Description of the experimental design: the same biological material is sampled at several times points across several omic layers indicated in different colors. Blank lines indicate potential missing values per time point or per feature in a given time point. **B**: Specific pre-processing and normalization are applied according to the type of data. **C**: Each molecule is modelled as a function of time by taking into account all the variability of the different biological replicates in a linear mixed model spline framework. **D**: The modelled trajectories across all omics layers are clustered using a multivariate integrative method.

## 3 Method

Our proposed approach includes pre-processing for microbiome data, spline modelization within a linear mixed model framework, and a multivariate analysis for clustering and data integration (Figure 2).

**Figure 2:**
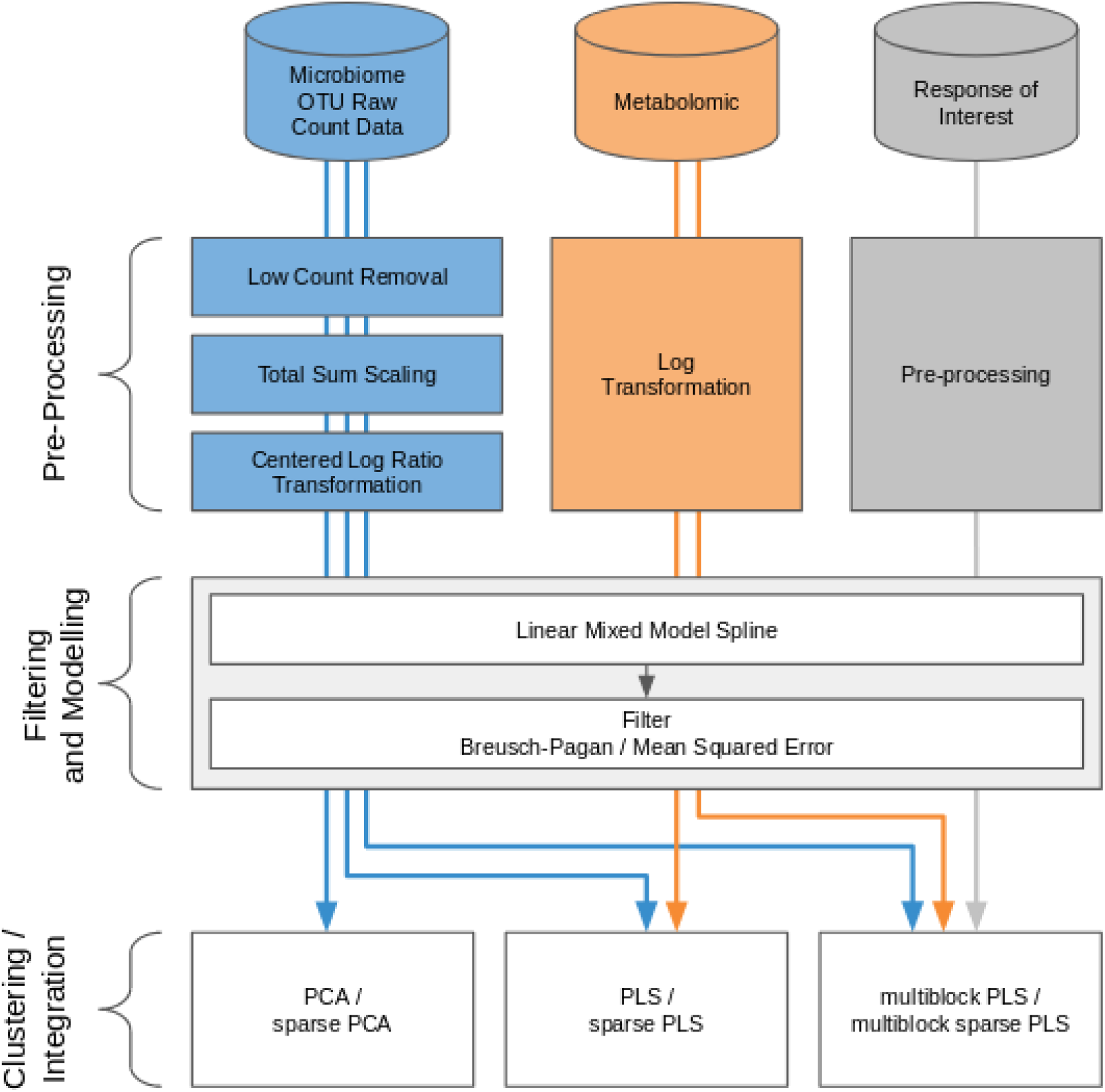
Workflow diagram for longitudinal integration of microbiota studies. We consider studies for the analysis of the microbiota through OTU (16S amplicon) or gene (whole genome shotgun) counts. This information can be complemented by additional information at the microbiota level, such as metabolic pathways measured with metabolomics, or information measured at a macroscopic level resulting from the aggregated actions of the microbiota.

**Figure 3:**
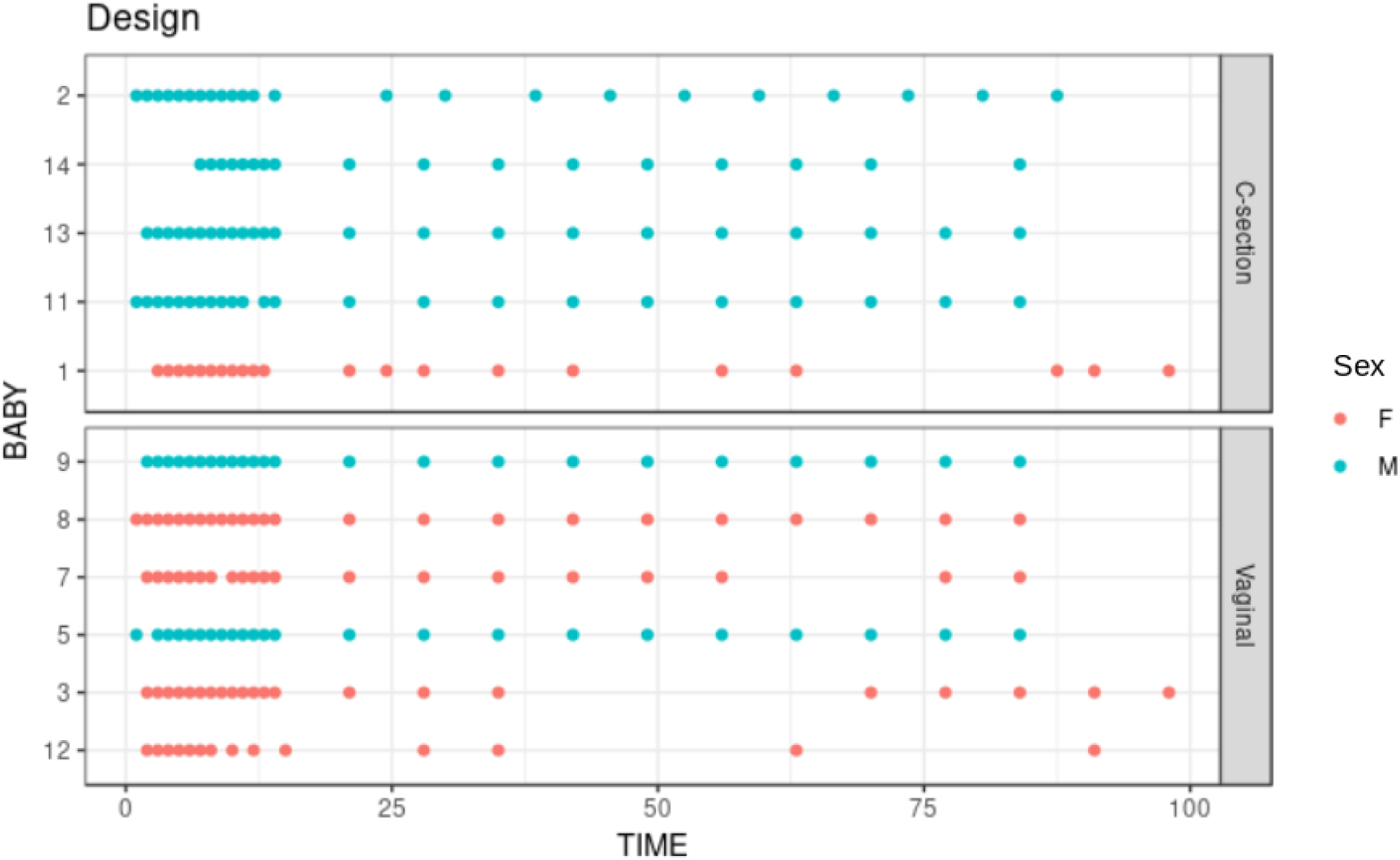
Infant gut microbiota development study: stool samples were collected from six male and five female babies over the course of 100 days. Samples were collected daily during days 0-14 and weekly thereon until day 100. Time is indicated on the x-axis in days. As delivery method is known to be a strong influence on gut microbiome colonization, the data are separated according to either C-section or vaginal birth.

**Figure 4:**
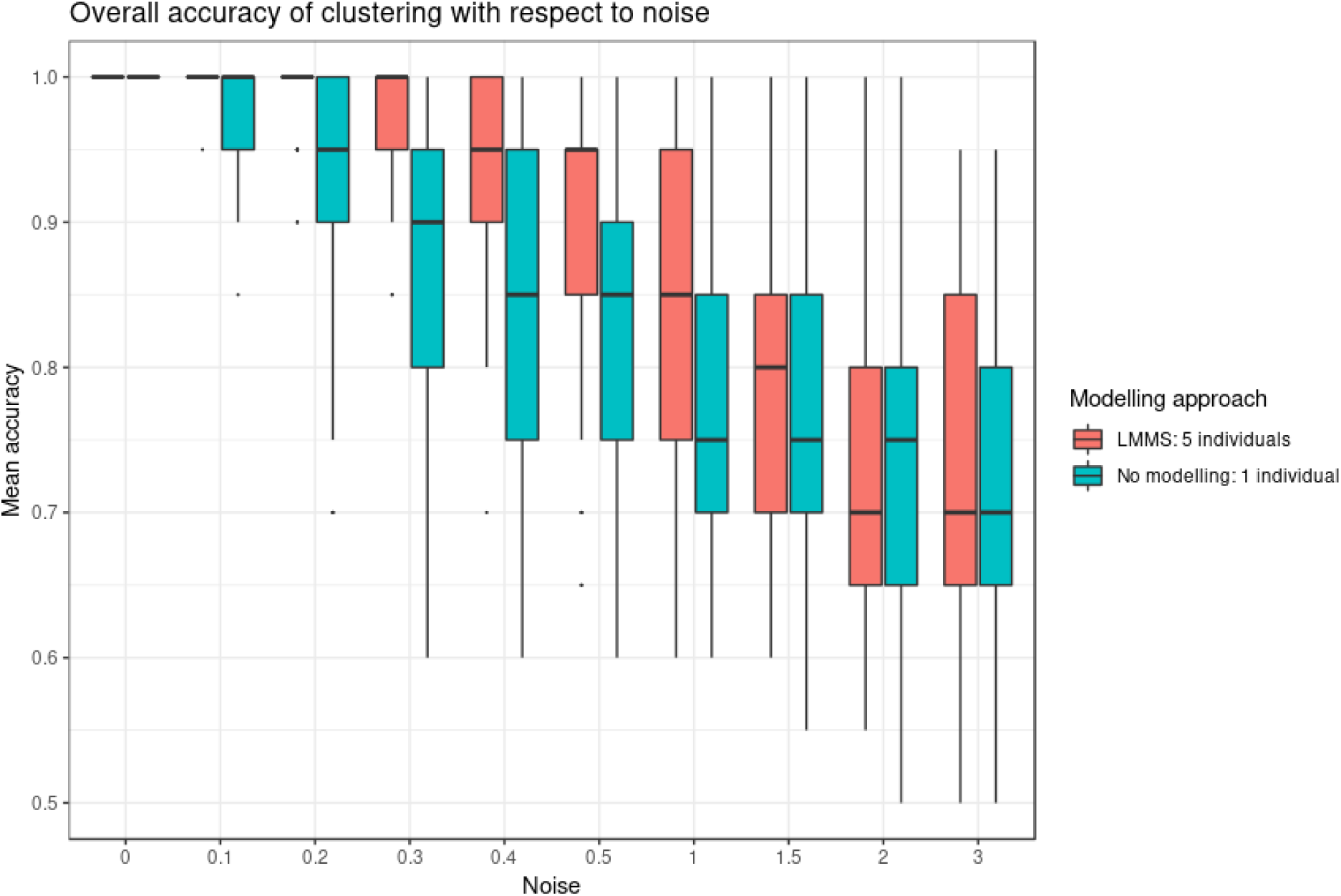
Simulation study: Overall accuracy of clustering with respect to noise. Twenty reference profiles, grouped into 4 clusters were used as a basis for simulation and each of the new simulated profiles were generated with random noise (described in Section 3.7.1). We compared two approaches: with LMMS modelling: 5 new profiles were generated per reference, and without modelling: only one profile was simulated per reference. We evaluated the ability of PCA clustering to correctly assign the simulated profiles in their respective reference clusters based on mean accuracy: without noise, both approaches lead to a perfect clustering, with Noise < 1, LMMS modelling acts as a denoising process with better performance than no modelling, and with a high level of noise ≥ 1 the performance of both approaches decrease.

**Figure 5:**
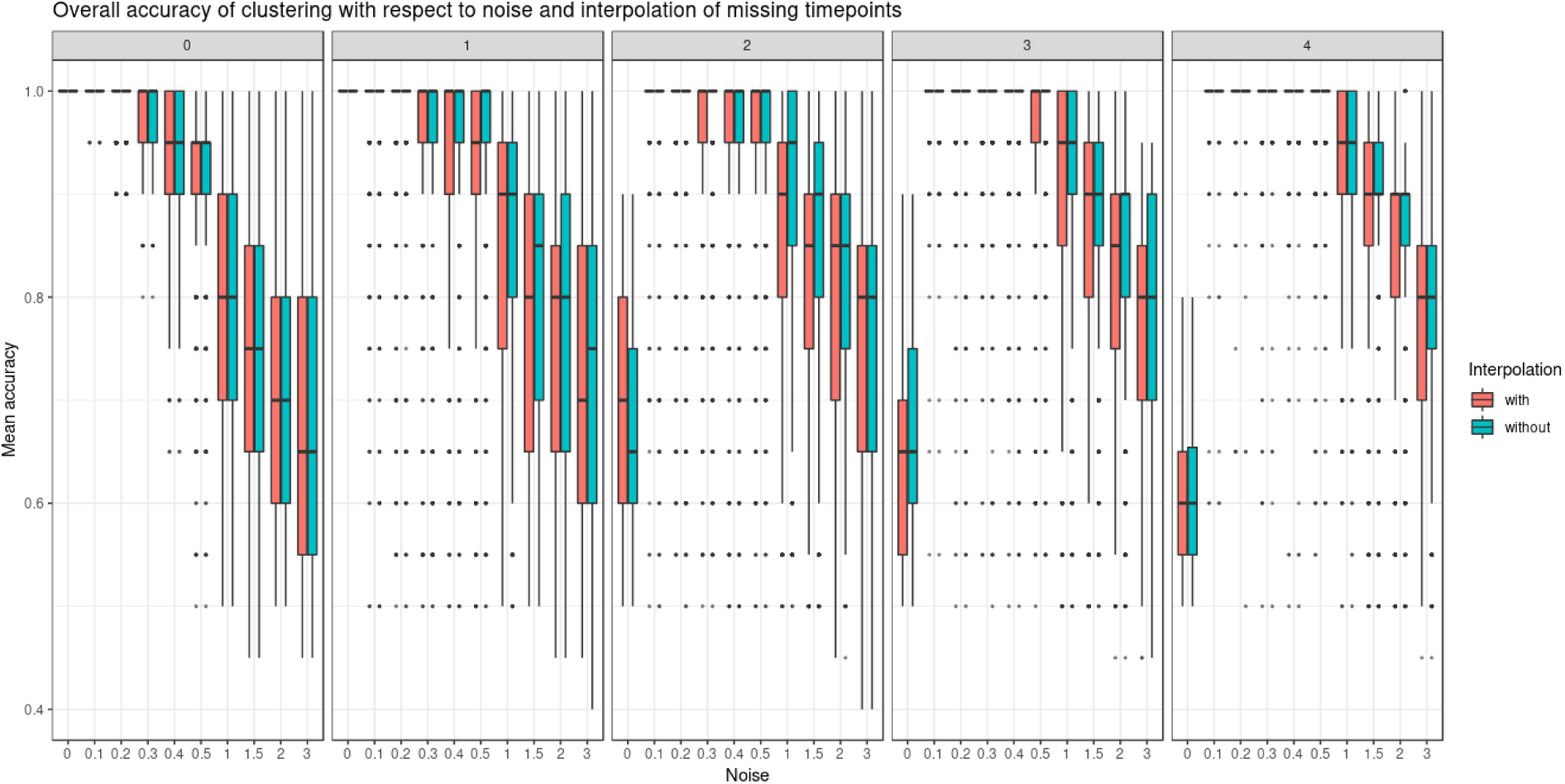
Simulated study: Overall accuracy of clustering when time points are missing. The simulation scheme is described in 3.7.1, however, here some time points were removed. We compared the ability of LMMS to interpolate missing time points. When there are no time points missing, both interpolated and non-interpolated approaches gave a similar performance. When the number of time points increases, the classification accuracy decreases. Without noise and with several timepoints removed, LMMS tended to model straight lines, resulting in poor clustering (see also Supplemental Figure S5).

**Figure 6:**
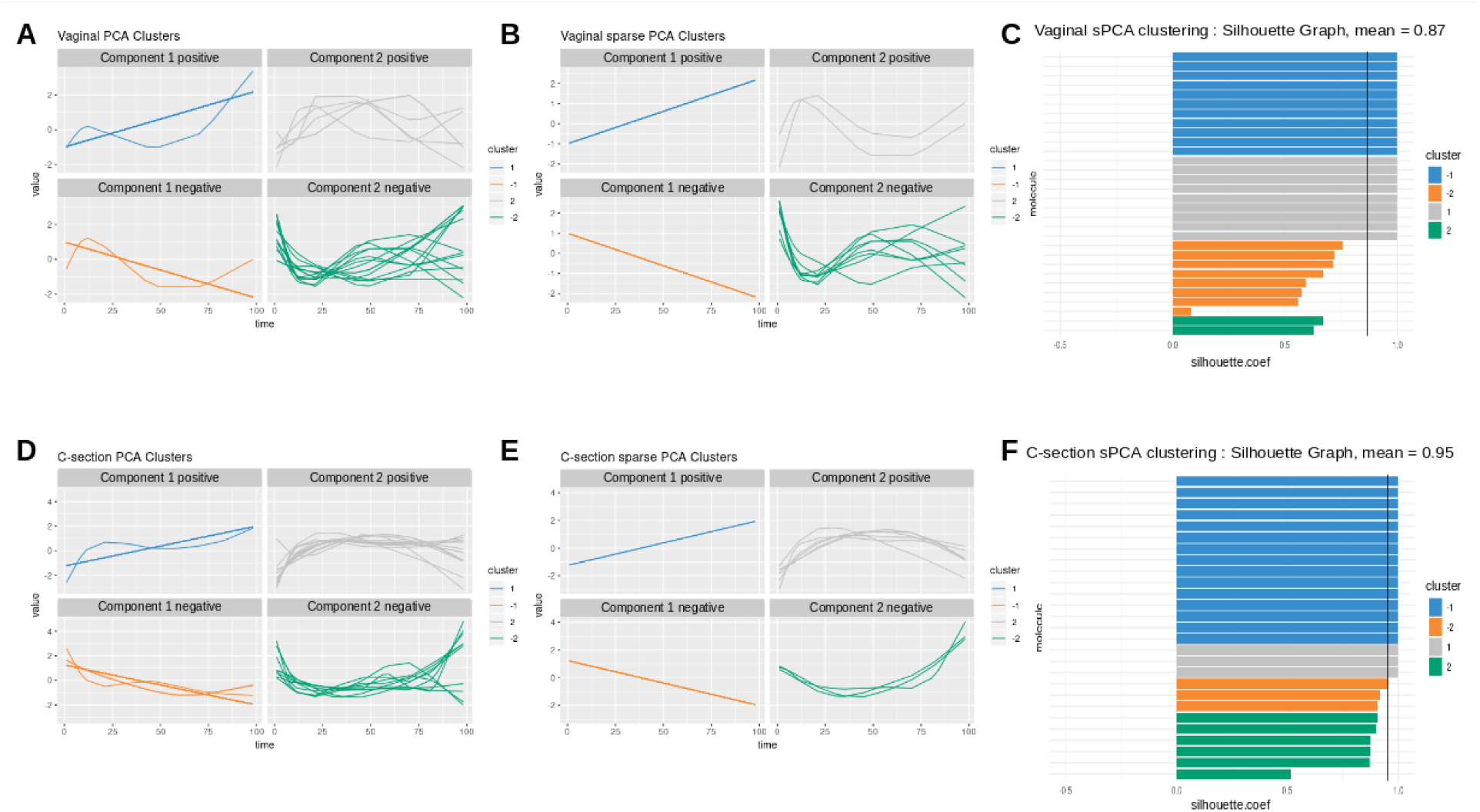
Infant gut microbiota development. Vaginal (first row) and C-section delivery babies (second row). (**A**) - (**B**), (**D**) - (**E**): OTU profiles clustered with either PCA or sPCA. Each line represents the abundance of an OTU across time. OTUs were clustered according to their contribution on each component for PCA or sPCA (that includes variable selection). The PCA clusters were further separated into profiles denoted ‘positive’ or ‘negative’ that refer to the sign of the loading vector from s/PCA. Profiles were scaled to improve visualization. (**C**) - (**F**): Silhouette profiles for each identified clustering. Each bar represents the silhouette coefficient of a particular OTU and colors represent assigned clusters. The average coefficient is represented by a vertical black line. A greater average silhouette coefficient means a better partitioning state. (**C**): vaginal delivery babies with sPCA (average silhouette coefficient = 0.87); (**F**): C-section delivery babies with sPCA (average silhouette coefficient = 0.95).

**Figure 7:**
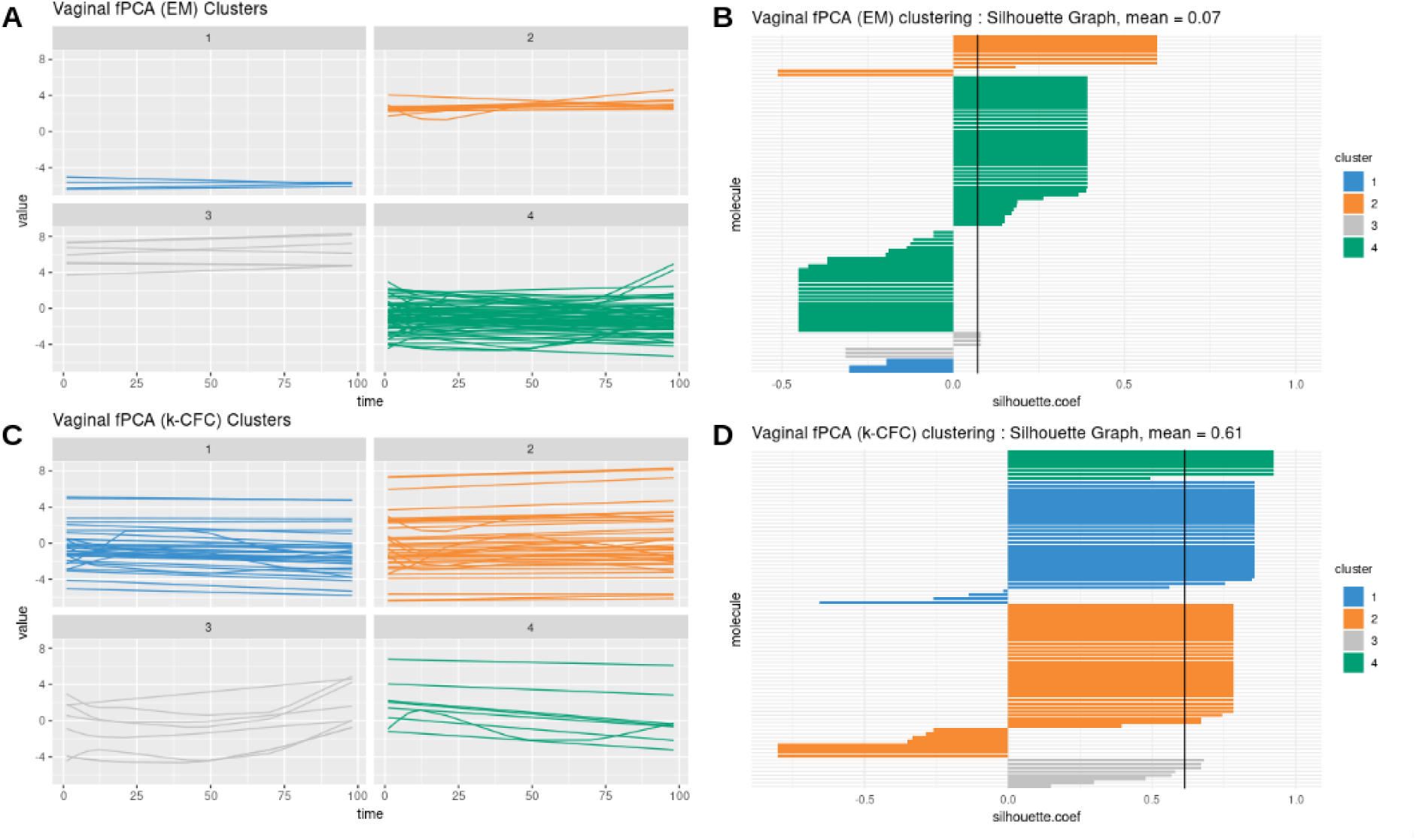
Infant gut microbiota development. fPCA Expectation-Maximization clustering (first row) and *k*-Center Functional Clustering (second row). (**A**) - (**C**): Vaginal OTU profiles clustered with either EM or *k*-CFC. Each line represents the relative abundance of a selected OTU across time. (**B**) - (**D**): Silhouette coefficients for each profile and each clustering. Each bar represents the silhouette coefficient of a particular OTU and colors represent assigned clusters. The average coefficient is represented by a vertical black line. The average silhouette coefficient was 0.07 for EM clustering and 0.61 for *k*-CFC clustering.

### 3.1 Pre-processing of microbiome data

We assume the data are in raw count formats resulting from bioinformatics pipelines such as QIIME (Caporaso *et al.*, 2010) or FROGS (Escudié *et al.*, 2017) for 16S amplicon data. Here we consider the OTU taxonomy level, but other levels can be considered, as well as other types of microbiome-derived data, such as whole genome shotgun sequencing. The data processing step is described in Lê Cao *et al.* (2016a) and consists of:

1. Low Count Removal: Only OTUs whose proportional counts exceeded 0.01% in at least one sample were considered for analysis. This step aims to counteract sequencing errors (Kunin *et al.*, 2010).
2. Total Sum Scaling (TSS) can be considered as a ‘normalisation’ process to account for uneven sequencing depth across samples. TSS divides each OTU count by the total number of counts in each individual sample but generates compositional data expressed as proportions.
3. Centered Log Ratio transformation (CLR) addresses in a practical way the compositionality issue, by projecting the data into a Euclidean space (Aitchison, 1982; Fernandes *et al.*, 2014; Gloor *et al.*, 2017). Given a vector *x* of *p* OTU counts for a given sample, CLR (eq. 1) is a log transformation of each element of the vector divided by its geometric mean *G*(*x*):

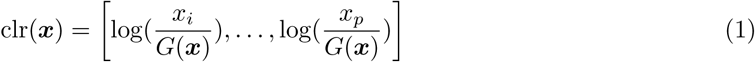

where

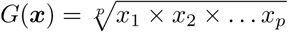

### 3.2 Time profile modelling

#### 3.2.1 Linear Mixed Model splines (LMMS)

The LMMS modelling approach proposed by Straube *et al.* (2015) takes into account between and within individual variability and irregular time sampling. LMMS is based on a linear mixed model representation of penalised splines (Durbán *et al.*, 2005) for different types of models. Through this flexible approach of serial fitting, LMMS avoids under- or over-smoothing. Briefly, four types of models are consecutively fitted in our framework on the TSS-CLR data:

1. A simple linear regression of taxa abundance on time, estimated via ordinary linear least squares - a straight line that assumes the response is not affected by individual variation.
2. A penalised spline proposed by Durbán *et al.* (2005) to model nonlinear response patterns.
3. A model that accounts for individual variation with the addition of a subject-specific random effect to the mean response in model (2).
4. An extension to model (3) that assumes individual deviations are straight lines, where individual-specific random intercepts and slopes are fitted.

All four models are described in Appendix A. Straube *et al.* 2015 showed that the proportion of profiles fitted with the different models increased in complexity with the organism considered. Different types of splines can be considered in models (2) - (4), including a cubic spline basis (Verbyla *et al.*, 1999), a penalised spline and a cubic penalised spline. A cubic spline basis uses all inner time points of the measured time interval as knots, and is appropriate when the number of time points is small (≤ 5), whereas the penalised spline and cubic penalised spline bases use the quantiles of the measured time interval as knots, see Ruppert (2002). In our case studies, we used penalised splines. The LMMS models are implemented in the R package `lmms` (Straube *et al.*, 2016).

#### 3.2.2 Prediction and interpolation

The fitted splines enable us to predict or interpolate time points that might be missing within the time interval (e.g. inconsistent time points between different types of data or covariates). Additionally, interpolation is useful in our multivariate analyses described below to smooth profiles, and when the number of time points is small (≤ 5). In the following section, we therefore consider data matrices ***X*** (*T* × *P*), where *T* is the number of (interpolated) time points and *P* the number of taxa. The individual dimension has thus been summarised through the spline fitting procedure, so that our original data matrix of size (*N* × *P* × *T*), where *N* is the number of biological samples, is now of size (*T* × *P*).

### 3.3 Filtering profiles after modelling

A simple linear regression model (1) might be the result of highly noisy data. To retain only the most meaningful profiles, the quality of these models was assessed with a Breusch-Pagan test to indicate whether the homoscedasticity assumption of each linear model was met (Breusch and Pagan, 1979). We also used a threshold based on the mean squared error (MSE) of the linear models, by only including profiles for which their MSE was below the maximum MSE of the more complex fitted models (1) - (4). The latter filter was only applied when a large number of linear models (1) were fitted and the Breusch-Pagan test was not considered stringent enough.

### 3.4 Clustering time profiles

#### 3.4.1 PCA and sparse PCA

Multivariate dimension reduction techniques such as Principal Component Analysis (PCA, Jolliffe 2005) and sparse PCA (Huang and Zheng, 2006) can be used to cluster taxa profiles. To do so, we consider as data input the *X* (*T* × *P*) spline fitted matrix. Let *t*_1_, *t*_2_, *…*, *t*_*H*_ denote the *H* principal components of length *T* and their associated *v*_1_, *v*_2_, *…*, *v*_*H*_ factors - or loading vectors, of length *P*. For a given PCA dimension *h*, we can extract a set of strongly correlated profiles by considering taxa with the top absolute coefficients in *v*_*h*_. Those profiles are linearly combined to define each component *t*_*h*_, and thus, explain similar information on a given component. Different clusters are therefore obtained on each dimension *h* of PCA, *h* = 1 *… H*. Each cluster *h* is then further separated into two sets of profiles which we denote as ‘positive’ or ‘negative’ based on the sign of the coefficients in the loading vectors (see for example Figures 6, 8 and 9).

**Figure 8:**
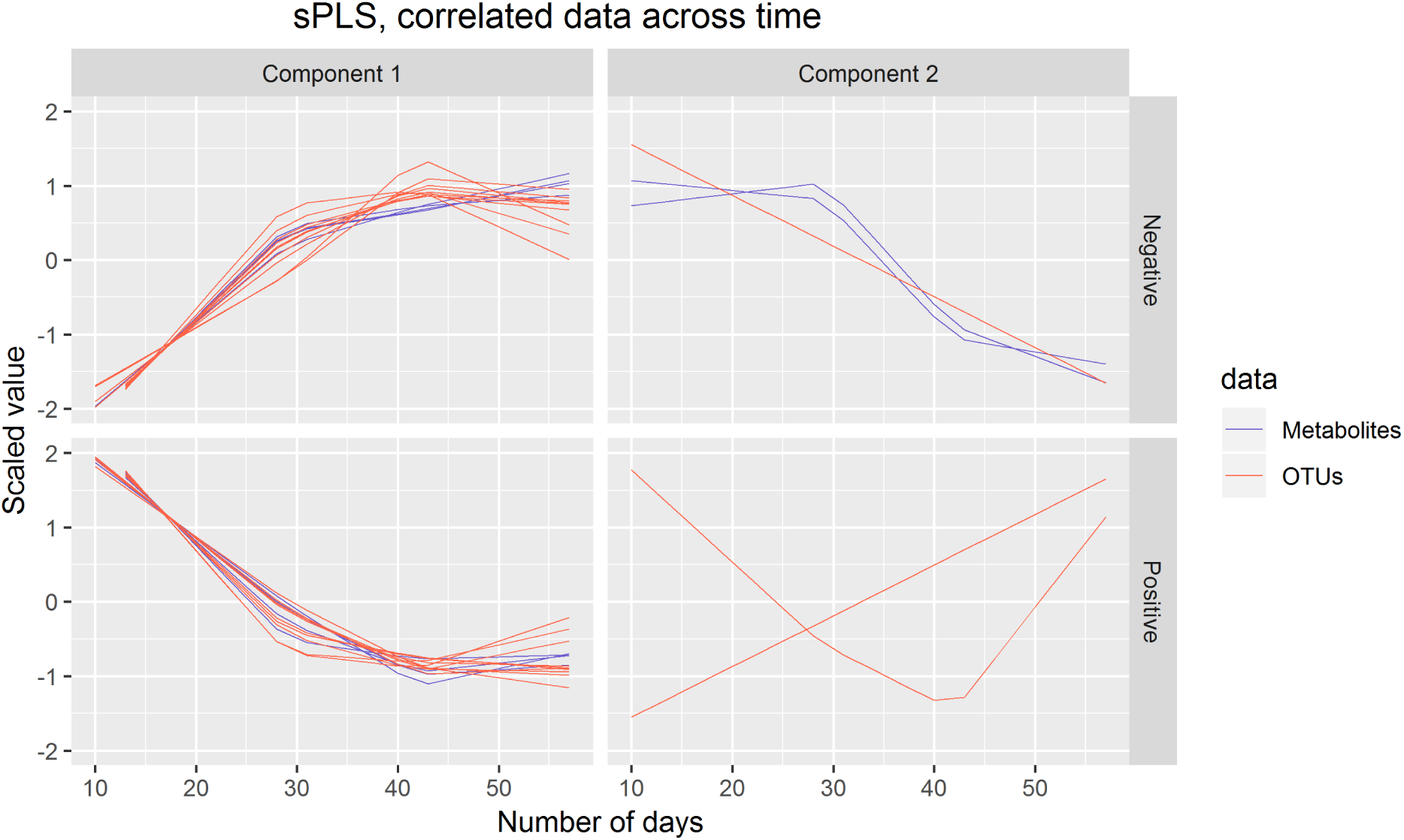
Waste degradation study: sPLS analysis identified subsets of associated OTUs and metabolites profiles. Each line represents the relative abundance of OTUs and metabolites selected by sPLS across time. OTUs and metabolites were clustered according to their contribution on each component. The clusters were further separated into profiles denoted ‘positive’ or ‘negative’ that refer to the sign of the loading vector from sPLS.

**Figure 9:**
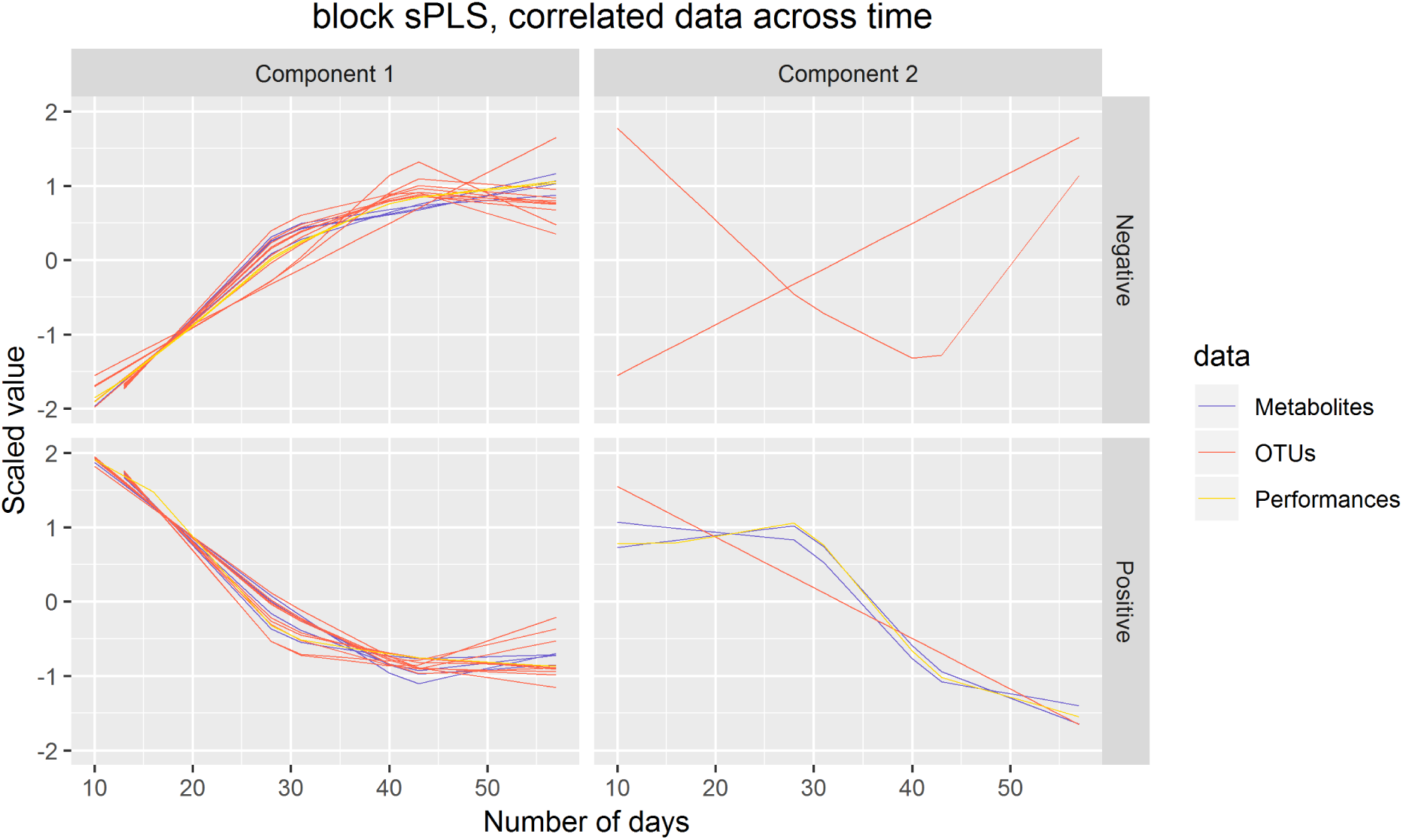
Waste degradation study: integration of OTUs, metabolites and performance measures with multiblock sPLS. Each line represents the relative abundance of OTUs, metabolites and performance measures selected by multiblock sPLS across time. OTUs, metabolites and performance measures were clustered according to their contribution on each component. The clusters were further separated into profiles denoted ‘positive’ or ‘negative’ that refer to the sign of the loading vector from multiblock sPLS.

A more formal approach can be used with sparse PCA. Sparse PCA includes 𝓁_1_ penalizations on the loading vectors to select variables that are key for defining each component, and are highly correlated within a component (see Huang and Zheng 2006 for more details).

#### 3.4.2 Choice of the number of clusters in PCA

We propose to use the average silhouette coefficient (Rousseeuw, 1987) to determine the optimal number of clusters, or dimensions *H*, in PCA. For a given identified cluster and observation *i*, the silhouette coefficient of *i* is defined as

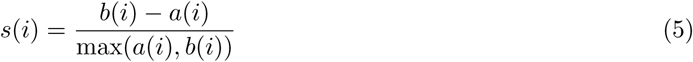

where *a*(*i*) is the average distance between observation *i* and all other observations within the same cluster, and *b*(*i*) is the average distance between observation *i* and all other observations in the nearest cluster. A silhouette score is obtained for each observation and averaged across all silhouette coefficients, ranging from −1 (poor) to 1 (good clustering).

We adapted the silhouette coefficient to choose the number of components or clusters in PCA and sPCA (i.e. 2 × *H* clusters), as well as the number of profiles to select for each cluster. Each observation in Eq. (5) now represents a fitted LMMS profile, and the distance between two profiles is calculated using the Spearman Correlation coefficient.

Within a given cluster, we calculate the silhouette coefficient of each LMMS profile and apply the following empirical rules for cluster assignation: a coefficient > 0.5 assigns the profile to the cluster, a value between 0 and 0.5 indicates an uncertain assignment as the profile can be assigned to one or two clusters, while a negative value indicates that the profile should not be assigned to this particular cluster.

To choose the appropriate number of profiles per sPCA component, we perform as follows: For each component, we set a grid of the number of profiles to be retained with sPCA and calculated the average silhouette coefficient per cluster (there are two clusters per component). The final number of profiles to select is arbitrarily set when we observe a sudden decrease in the average silhouette coefficient (see Figure 7).

#### 3.4.3 Comparison with Functional Principal Component Analysis (fPCA)

fPCA has been widely used to cluster longitudinal data by decomposing data matrices into temporal variation models (Hyndman and Ullah, 2007) and has been used in several biological applications (Yao *et al.*, 2005; Silverman *et al.*, 1996). fPCA first models longitudinal profiles into a finite basis of functions, then clusters the longitudinal profiles using the basis expansion coefficients of the fPCA scores. fPCA requires the user to choose the number of clusters and the number of components - based on AIC, BIC, or percentage of total explained variance, the approach to estimate the fPCA scores - based on conditional expectation or numerical integration, and to cluster the profiles. We used the ‘fdapace’ R package that includes two types of clustering methods, based on ‘model-based clustering of finite mixture Gaussian distribution (‘EMCluster’) or k-means algorithm based on the fPCA scores.

### 3.5 Evaluation

#### 3.5.1 Clustering

We can assess the quality of clustering with internal measures such as compactness (Dunn, Rand indices, and Jaccard index) or cluster separation. For the latter case, the silhouette coefficient is recognized as an informative criterion Wang *et al.* (2009), and can be used to compare several clustering results based on the same data. Thus we used this criterion to assess different methods (PCA, sPCA and fPCA), or to assess the same method with different parameters, for example to identify the appropriate number of clusters as we described in 3.4.2. The best clustering approach yields the highest silhouette coefficient.

#### 3.5.2 Measure of association for compositional data

Compositional data arise from any biological measurement made based on relative abundance (Gloor *et al.*, 2017; Lovell *et al.*, 2015). Microbiome data in particular are compositional for several reasons, including biological, technical and computational. Thus, interpretation based on correlations between profiles must be made with caution as it is highly likely to be spurious. Proportional distances has been proposed as an alternative to measure association. The compositional data analysis field is an active field of research but methods are critically lacking for longitudinal data. Here we adopt a practical and post-hoc approach to evaluate pairwise associations of microbial and omics profiles once they have been assigned to their clusters. We used the proportionality distance *φ*_*s*_ proposed by Lovell *et al.* (2015) and implemented in the ‘propr’ R package (Quinn *et al.*, 2017). For two LMMS profiles *x*_*i*_ and *x*_*j*_, we define the pairwise proportionality distance as

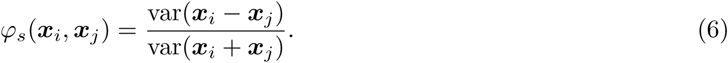

A small value indicates that, in proportion, the pair of profiles are strongly associated. We calculated the distance *φ*_*s*_ on the log-transformed LMMS modelled profiles within each identified cluster to exclude potentially spurious correlations and further guide the interpretation of the results. In addition, to evaluate the quality of our clustering approach, we compared the pairwise distances of the profiles within a particular cluster and profiles outside the cluster.

### 3.6 Integration

#### 3.6.1 Multiblock PLS methods

To integrate multiple datasets (also called *blocks*) measured on the same biological samples, we used multivariate methods based on Projection to Latent Structures (PLS) methods (Wold, 1975), which we broadly term *multiblock PLS* approaches. For example, we can consider Generalised Canonical Correlation Analysis (GCCA, Tenenhaus and Tenenhaus 2011; Tenenhaus *et al.* 2014), which, contrary to what its name suggests, generalises PLS for the integration of more than two datasets. Recently, we have developed the DIABLO method to discriminate different phenotypic groups in a supervised framework (Singh *et al.*, 2019). In the context of this study however, we present the sparse GCCA in an unsupervised framework, where input datasets are spline-fitted matrices.

We denote *Q* data sets *X*^(1)^(*T* × *P*_1_), *X*^(2)^(*T* × *P*_2_), …, *X*^(*Q*)^(*T* × *P*_*Q*_) measuring the expression levels of *P*_*q*_ variables of different types (taxa, ‘omics, continuous response of interest), modelled on *T* (interpolated) time points, *q* = 1, *…*, *Q*. GCCA solves for each component *h* = 1, *…*, *H*:

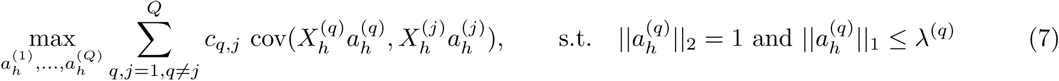

where *λ*^(*q*)^ is the 𝓁_1_ penalization parameter, 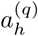 is the loading vector on component *h* associated with the residual (deflated) matrix 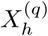 of the data set *X*^(*q*)^, and *C* = {*c*_*q,j*_}_*q,j*_ is the design matrix. *C* is a *Q* × *Q* matrix that specifies whether datasets should be correlated and includes values between zero (datasets are not connected) and one (datasets are fully connected). Thus, we can choose to take into account specific pairwise covariances by setting the design matrix (see Rohart *et al.* 2017 for implementation and usage) and model a particular association between pairs of datasets, as expected from prior biological knowledge or experimental design. In our integrative case study, we used sparse PLS, a special case of Eq. (7) to integrate microbiome and metabolomic data, as well as sparse multiblock PLS to also integrate variables of interest. Both methods were used with a fully connected design.

The multiblock sparse PLS method was implemented in the `mixOmics` R package where the *𝓁*_1_ penalization parameter is replaced by the number of variables to select, using a soft-thresholding approach (see more details in Rohart *et al.* 2017).

#### 3.6.2 Parameters tuning

The integrative methods require choosing the number of components *H*, defined as 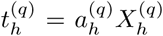 with the notations from Section 3.6.1, and number of profiles to select on each PLS component and in each dataset. We generalized the approach described in Section 3.4.2 using the silhouette coefficient based on a grid of parameters for each dataset and each component.

### 3.7 Simulation and case studies

#### 3.7.1 Simulation study description

A simulation study was conducted to evaluate the clustering performance of multivariate projection-based methods such as PCA, and the ability to interpolate time points in LMMS.

Twenty reference time profiles were generated on 9 equally spaced time points and assigned to 4 clusters (5 profiles each). These ground truth profiles were then used to simulate new profiles. We generated 500 simulated datasets.

*Clustering performance.* We first compared profiles simulated then modelled with or without LMMS:

A. For each of the reference profiles, 5 new profiles (corresponding to 5 individuals) were sampled to reflect some inter-individual variability as follows: Let *x* be the observation vector for a reference profile *r, r* = 1 *…* 20, for each time point *t* (*t* = 1, *…*, 9), 5 measurements were randomly simulated from a Gaussian distribution with parameters *µ* = *x*_*t,r*_ and *σ*^2^, where *σ* = {0, 0.1, 0.2, 0.3, 0.4, 0.5, 1, 1.5, 2, 3} to vary the level of noise. This noise level was representative of the data described below in Section gut microbiota development. The profiles from the 5 individuals were then modelled with LMMS (section 3.2.1, resulting in 500 matrices of size (9 × 20).
B. For each of the reference profiles, one new profile was simulated as described in step A, but no LMMS modelling step was performed, resulting in 500 matrices of size (9 × 20).

Clustering was obtained with PCA (see Section 3.4.1) and compared to the reference cluster assignments in a confusion matrix. The clustering was evaluated by calculating the accuracy of assignment 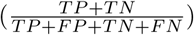 from the confusion matrix, where for a given cluster, TP (true positive) is the number of profiles correctly assigned in the cluster, FN (false negative) is the number of profiles that have been wrongly assigned to another cluster, TN (true negative) is the number of profiles correctly assigned to another cluster and FP (false positive) is the number of profiles incorrectly assigned to this cluster. Besides accuracy, we also calculated the Rand index (Rand, 1971) as a similarity metric to the clustering performance of PCA. The clustering results from fPCA were poor, even for a low level of noise (Suppl. Figure S6), thus fPCA was not compared against PCA.

*Interpolation of missing time points.*

To evaluate the ability of LMMS to predict the value of a missing time point for a given feature over time, we randomly removed 0 to 4 measurement points in the simulated datasets described above in step A. We compared the PCA clustering performance with or without LMMS interpolation.

#### 3.7.2 Infant gut microbiota development

The gastrointestinal microbiome of 14 babies during the first year of life was studied by Palmer *et al.* (2007). The authors collected an average of 26 stool samples from healthy full-term infants. As infants quickly reach an adult-like microbiota composition, we focused our analyses on the first 100 days of life. Infants who received an antibiotic treatment during that period were removed from the analysis, as antibiotics can drastically alter microbiome composition (Dudek-Wicher *et al.*, 2018).

The dataset we analysed included 21 time points on average for 11 selected infants (Figure 3). Samples were collected daily during days 0-14 and weekly after the second week. We separated our analyses based on the delivery mode (C-section or vaginal), as this is known to have a strong impact on gut microbiota colonization patterns and diversity in early life (Rutayisire *et al.* (2016)). The purpose of our statistical analysis was to identify a bacterial signature that describes the dynamics of a baby’s microbial gut development in the first days of life, as well as compare differences in signatures between babies born by vaginal delivery or by C-section. As this study is single omics, we applied our framework depicted in Figure 2 with sPCA.

#### 3.7.3 Waste degradation study

Anaerobic digestion (AD) is a highly relevant microbial process to convert waste into valuable biogas. It involves a complex microbiome that is responsible for the progressive degradation of molecules into methane and carbon dioxide. In this study, AD’s biowaste was monitored across time (more than 150 days) in three lab-scale bioreactors as described in (Poirier *et al.*, 2016).

We focused our analysis on days 9 to 57, that correspond to the most intense biogas production. Degradation performance was monitored through 4 parameters: methane and carbon dioxide production (16 time points) and the accumulation of acetic and propionic acid in the bioreactors (5 time points). Microbial dynamics were profiled with 16S RNA gene metabarcoding as described in Poirier *et al.* (2016), and included 4 time points and 90 OTUs. A metabolomic assay was conducted on the same biological samples at 4 time points with gas chromatography coupled to mass spectrometry GC-MS after solid phase extraction to monitor substrates degradation (Limam *et al.*, 2010). The XCMS R package (version 1.52.0) was used to process the raw metabolomics data (Smith *et al.*, 2006). GC-MS analyses focused on 20 peaks of interest identified by the National Institute of Standards and Technology database. Data were then log-transformed. The purpose of the study was to investigate the relationship between biowaste degradation performance, microbial and metabolomic dynamics across time. The aim of our statistical analysis was to identify highly associated multi-omic signatures characterizing waste degradation dynamics in the 3 bioreactors. This study involves the integration of two omics datasets and degradation performance measures, thus we applied sPLS and multiblock sPLS, as shown in our workflow in Figure 2.

## 4 Results

### 4.1 Simulation study

#### 4.1.1 Clustering performance

Figure 4 shows the clustering performance of PCA with an increasing amount of noise in the simulated profiles. Unsurprisingly, PCA gave optimal clustering performance when noise was absent, with or without profile modelling to take into account individual variability. When noise increased, PCA performed better with modelling, which acts as a denoising process. Finally, a high level of noise showed the limitation of the modelling approach, as similar clustering results were obtained with or without LMMS modelling. However, the PCA clustering performance was still very good, with a mean accuracy of 0.7 when the level of noise was maximum.

#### 4.1.2 Interpolation of missing time points

We evaluated the ability of LMMS to interpolate an increasing number of missing time points (up to four). Interpolation is important in our framework as it allows the estimation of evenly spaced time points as well as time points that may be missing in one data set but not in the other (e.g biowaste degradation study described in 3.7.3). Interpolation did not seem to affect the clustering performance of PCA (Supplementary Figures 5 and S1). Rather, the level of noise had the largest impact on clustering: the mean accuracy was close to 1 when the noise was non existent, but decreased as the number of missing time points and noise increased. In the latter scenarios, LMMS interpolation seemed to give, on average, better clustering than without interpolation. When the number of missing time points increased, we observed a better classification accuracy with noise compared to no noise. This can be explained by the LMMS modelling of straight lines in the latter case that led to poor clustering (Supplementary Figure S5).

### 4.2 Clustering time profiles: Infant gut microbiota development study

#### 4.2.1 Pre-processing and modelling

A total of 2,149 taxa were identified in the raw data (Table 1). After the pre-processing steps illustrated in Figure 2, a smaller number of OTUs were found in faecal samples of babies born by C-section than vaginal delivery. Similarly, a simple linear regression model showed a smaller proportion of OTUs in babies born via C-section (73%) than vaginal delivery (81%), and this was also observed after the filtering step (Table 1).

**Table 1:** Infant gut microbiota development study: Number of OTUs identified, and linear model types fitted according to delivery mode.

|  |  | C-section | Vaginal |
| --- | --- | --- | --- |
| Identified OTUs |  | 2,149 | 2,149 |
| # OTUs after pre-processing |  | 107 | 117 |
| Linear model types | (1) | 78 | 95 |
|  | (2) | 29 | 22 |
| Linear model types after filtering | 1 | 42 | 68 |
|  | 2 | 29 | 22 |

#### 4.2.2 Comparison of PCA and functional PCA

According to our tuning criteria, we obtained four clusters with PCA (i.e. two components). We therefore set the same number of clusters in fPCA for comparative purposes. PCA clustering outperformed fPCA for each delivery mode dataset that was analyzed (see Table 2). The resulting fPCA clustering is displayed in Figure 7 for babies born via vaginal delivery. We found that the EM approach in fPCA tended to cluster a larger number of uncorrelated OTUs compared to the *k*-CFC approach (average silhouette coefficient = 0.07 for EM and 0.61 for *k*-CFC).

**Table 2:** Infant gut microbiota development study: Average silhouette coefficient according to clustering method

|  | PCA | sPCA | fPCA (*k*-CFC) | fPCA(EM) |
| --- | --- | --- | --- | --- |
| Vaginal | 0.84 | 0.95 | 0.61 | 0.07 |
| C-section | 0.87 | 0.86 | 0.69 | 0.35 |

#### 4.2.3 Clusters of profiles

We used sPCA to select key OTU profiles for each cluster. This step is essential for discarding profiles that are distant from the average cluster profile and thus not informative. As expected, we observed an overall increase in the silhouette average coefficient for the sPCA clustering compared to PCA, indicating a better clustering capability (see Table 2). According to the silhouette average coefficient, vaginal delivery showed the best partitioning for PCA clustering (0.87, Table 2). Cluster 1 (denoted ‘component 1 positive’ in Figure 6 **A**) showed a relative increase in abundance of species, including some that are characteristic of a healthy “adult-like” gut microbiome composition such as the clade *Bacteroidetes* (Thursby and Juge, 2017). The proportionality distance within cluster 1 was low (Supplementary Table S1), with a strong association between *Bacteroides* and *Fusobacteria* (*φ*_*s*_ = 0.04), as well as between *Actinobacter* with *Bacteroides* (*φ*_*s*_ = 0.02) and *Fusobacteria*(*φ*_*s*_ = 0.09). According to this distance, there might have been a spurious correlation identified between the genus *Bacteroides* and an environmental uncultured bacterium *(clone HuCA36)* (*φ*_*s*_ = 14.81), see Supplementary Table S4. In cluster 2 (‘component 1 negative’), relative profile abundance tended to decrease and corresponded to genera found in vaginal and skin microbiota, such as *Lactobacillus* and *Propionibacterium* (Grice and Segre, 2011; Bing *et al.*, 2012). According to the proportionality distance, *Propionibacterium* and *Lactobacillus* were highly associated (*φ*_*s*_ = 0.29) as well as with *Campylobacter* (*φ*_*s*_ = 0.39) (see Supplementary Table S4). Clusters 3 and 4 (denoted ‘component 2 positive and negative’) highlighted taxa profiles with negative association.

Thus, with this preliminary PCA analysis, we were able to rebuild a partial history behind the development of the gut microbiota. Vaginal species that initially colonized in the gut progressively disappeared to enable species that characterize adult gut microbiota.

For babies born by C-section, 4 clusters were identified by PCA (Fig. 6 **D**). The median values of the proportionality distance within the different clusters were significantly lower than between the selected OTUs in the clusters and all the other OTUs (Supplementary Table S2). For example, the median value within cluster 1 was 0.11 compared to 1.36 outside the cluster. Clusters 1 and 2 (‘component 1 positive and negative’) displayed either an increase or decrease in relative abundance. However none of the cluster 2 species are known to characterize, or were found in, vaginal delivery, suggesting that the infant gut was first colonized by the operating room microbes as already demonstrated by Shin *et al.* (2015). Cluster 3 (‘component 2 positive’) revealed transitory states of increase then decrease of relative abundance profiles, while cluster 4 (‘component 2 negative’) showed the inverse trend.

When comparing the dynamics of the 2 delivery methods, we found a higher diversity in the intestinal microbiota of babies born vaginally (117 modelled profiles) than than by C-section (107). For vaginal delivery, the modelling step identified a larger proportion of straight lines, which may indicated a greater inter-individual variability compared to C-section delivery. The clusters denoted ‘component 1 positive’ in both delivery modes showed an increased relative abundance over time, with 32 OTUs assigned to this cluster in vaginally born babies, compared to 11 in C-section (Table 3). Despite the relatively sterile environment of the operating room, it was surprising to observe similar number of OTUs in cluster ‘component 1 negative’ for both types of delivery mode (vaginal: 38, C-section: 35), as we would have expected to identify a larger number of opportunistic microorganisms colonizing babies born vaginally(e.g. *P. Acnes, Campylobacter*). These include species found on the surface of the skin and in the vaginal flora. However, for babies born by C-section we observed a large number of microorganisms from various origins (e.g. *Staphylococcus, Rickettsia, Rhodobacter*).

**Table 3:** Infant gut microbiota development study: Number of OTUs per cluster identified with PCA clustering and OTUs selected in brackets with sparce PCA.

|  | C-section | Vaginal |
| --- | --- | --- |
| Cluster 1 (comp 1 positive) | 11 (3) | 32 (9) |
| Cluster 2 (comp 1 negative) | 35 (15) | 38 (11) |
| Cluster 3 (comp 2 positive) | 15 (6) | 6 (2) |
| Cluster 4 (comp 2 negative) | 10 (3) | 14 (8) |

In summary, sparse PCA clustering of LMMS modelled profiles enabled the identification of groups of microorganisms with relative increased abundance over time. These microorganisms are characteristics of an adult gut microbiota. We also identified groups of opportunistic microorganisms with a decreasing relative abundance over time. We also found that, during the first year of life, gut microbiota was more diverse for babies born by vaginal than C-section delivery.

### 4.3 Clustering omics: waste degradation study

#### 4.3.1 Pre-processing and modelling

A total of ninety OTUs were identified in the 12 samples of the initial dataset (Table 4). After preprocessing, 51 OTUs were retained. Approximately 60% (resp. 50%) of the OTUs (resp. metabolites) were fitted with linear regression models (1) and 40% (resp. 50%) were modelled by more complex splines models (2) - (4). All performance measures were also modelled by splines. During the filtering step, 7 OTUs and 4 metabolites that were fitted with linear regression models were discarded. The small number of profiles that were filtered out (described in Section 3.3) indicated that the variability between the 3 bioreactors was relatively low.

**Table 4:** Waste degradation study: OTUs, metabolites and performance modelling and filtering in the bioreactor study. Only OTU data were pre-processed.

| Type of features |  | OTUs | Metabolites | Performance |
| --- | --- | --- | --- | --- |
| Number of features |  | 90 | 20 | 4 |
| # features after pre-processing |  | 51 | NA | NA |
| Linear model types | (1) | 30 | 10 | 0 |
|  | (2) | 19 | 0 | 2 |
|  | (3) | 2 | 4 | 0 |
|  | (4) | 0 | 6 | 2 |
| Linear model types after filtering | (1) | 24 | 6 | 0 |
|  | (2) | 19 | 0 | 2 |
|  | (3) | 2 | 4 | 0 |
|  | (4) | 0 | 6 | 2 |

#### 4.3.2 sPCA on concatenated datasets

As a first and naive attempt to jointly analyze microbial, metabolomic and performance measures, all three datasets were concatenated then analyzed with sPCA. Only a very small number of profiles from the different datasets were selected. This small selection is likely due to the high variability in each data type. Selected variables included mainly OTUs and performance measures. These were assigned to four clusters and included respectively 1, 3, 2 and 3 OTUs with 0, 1, 2 and 0 metabolites and 2, 0, 1 and 0 performance measures. The average silhouette coefficient was 0.744, a potentially sub-optimal clustering compared to our analyses presented in the next Section. This preliminary investigation highlighted the limitation of sPCA to identify a sufficient number of associated profiles from disparate sources.

#### 4.3.3 Microbiome - metabolomic integration with sPLS

The results from the sPLS analysis are shown in Figure 8. Four clusters of variables were identified, and the average silhouette coefficient of 0.954 confirmed that sPLS led to better clustering of the different types of profiles than sPCA. The proportionality distances of the profiles within each cluster are presented in Table 5 and in Supplementary Figure S2. Their low values indicated strong associations between profiles within each cluster, compared to any association outside each of the clusters. The first cluster (denoted ‘component 1 negative’) included 10 OTUs and 4 metabolite variables, and showed increasing relative abundance until a plateau was reached at approximately 40 days. Median value of the proportionality distance within the cluster was 0.42, compared to 1.11 between the variables selected in the cluster and all the other variables, indicating strong associations within each cluster. The OTUs were microorganisms often recovered during anaerobic digestion of biowaste, such as methanogenic archaea of *Methanosarcina* genus or bacteria of *Clostridiales, Acholeplasmatales*, and *Anaerolineales* orders. These were reported as being involved in the different steps of anaerobic digestion (Poirier *et al.*, 2016). Their relative abundance increased while biowaste was degraded, until there was no more biowaste available in the bioreactor.

**Table 5:** Waste degradation study: Proportionality distance for clusters identified with sparse PLS. The median distance between all pairs of profiles, within cluster and with the entire background set (outside a given cluster) is reported. A Wilcoxon test p-value assesses the difference between the medians.

| cluster | median within cluster | median outside cluster | Wilcoxon test P-value |
| --- | --- | --- | --- |
| 1 (comp 1 positive) | 0.43 | 1.37 | $9.40 \times 10^{-57}$ |
| -1 (comp 1 negative) | 0.42 | 1.11 | $1.76 \times 10^{-28}$ |
| 2 (comp 2 positive) | 0.29 | 0.97 | $5.71 \times 10^{-24}$ |
| -2 (comp 2 negative) | 0.01 | 0.87 | $2.82 \times 10^{-13}$ |

From the proportionality distances, we found that their abundance across time was, in proportion, similar, indicating a synchronized role during this biological process. In particular, of all the proportionality distances between the profiles of archaea of *Methanosarcina* genus and bacteria of *Clostridiales* order, the *Syntrophomonadaceae* family was the lowest which made sense as these microorganisms have already been reported as syntrophs (Liu *et al.*, 2011), see Supplementary Material S5.

Their abundance was also highly associated, in proportion, to the intensity of various metabolites produced during the AD process, such as benzoic acid that is formed during the degradation of phenolic compounds (Hoyos-Hernandez *et al.*, 2014), or phytanic acid, known to be produced during the fermentation of plant materials in the ruminant gut (Watkins *et al.*, 2010), as well as indole-2-carboxylic acid. Thus, the identified microorganisms were likely responsible for the production of these compounds. Cluster 2 (component 1 positive) included 10 OTUs and 4 metabolites. The median value of the proportionality distance within the cluster was also very low compared to the proportionality distance outside the cluster (0.29 and 0.97, Table 5). Profiles of cluster 2 were negatively correlated to Cluster 1, and their relative abundance decreased with time. OTUs mainly belonged to the *Bacteroidales* order. They were present in the initial inoculum but did not survive in this experiment, as the operating conditions or the substrate were not optimal for their growth, as observed in other studies (Madigou *et al.*, 2019). Consequently, their relative abundance progressively decreased over time. Metabolites identified in Cluster 2 were present in the biowaste and were degraded during the experiment. They included fatty acids (decanoic and tetradecanoic acids) that can be found in oil, or 3-(3-Hydroxyphenyl)propionic acid, arising from the digestion of aromatic amino-acids or breakdown product of lignin or other plant-derived phenylpropanoids. As their profile was negatively correlated to those from cluster 1, it is likely that these metabolites were consumed by OTUs assigned to cluster 1 (Torres *et al.*, 2003). Cluster 3 (component 2 negative) included 1 OTU and 5 metabolites. Profiles relative abundance decreased slowly with time until reaching a stable abundance after 20 days. One OTU of *Clostridiales* order appeared to have been out-competed by other OTUs or phase active only during the first days of the degradation, which corresponds to the degradation of complex biopolymers contained in biowaste (Poirier *et al.*, 2016). Among the metabolites of this cluster, Hydrocinnamic and 3,4-Dihydroxyhydrocinnamic acids are commonly found in plant biomass and its residues (Boerjan *et al.*, 2003). Their molecular structure may have contributed to their slower degradation compared to other molecules, which may explain their stable abundance in the digesters until day 30. Finally, Cluster 4 (component 2 positive) included 11 OTUs and 3 metabolites with slow relative abundance increase. OTUs from this group were very varied with 8 orders represented. They may have had slower growth rates than OTUs of cluster 1 or were possibly involved in the degradation of molecules from cluster 3. Their abundance may also have had a slow increase as they fed on specific molecules that are only formed during the digestion process. Metabolites included N-Acetylanthranilic acid and Dehydroabietic acid that were likely produced by microorganisms and accumulated during the anaerobic digestion process, suggesting they could not be metabolised by other microorganisms.

#### 4.3.4 Microbiome, metabolomic and performance data integration with multiblock sPLS

Figure 9 illustrates the results from the integration of the three datasets, where the performance data are considered as the response of interest. Similar to the sPLS analysis, block sPLS assigned profiles to four clusters, with an average silhouette coefficient Additional Requirementsof 0.909. The proportionality distances are summarized in Figure 10 and in Table S3 and show a greater level of association between profiles within each cluster, compared to the associations with all other profiles outside the cluster (see Suppl Figure S4 per omic variable).

**Figure 10:**
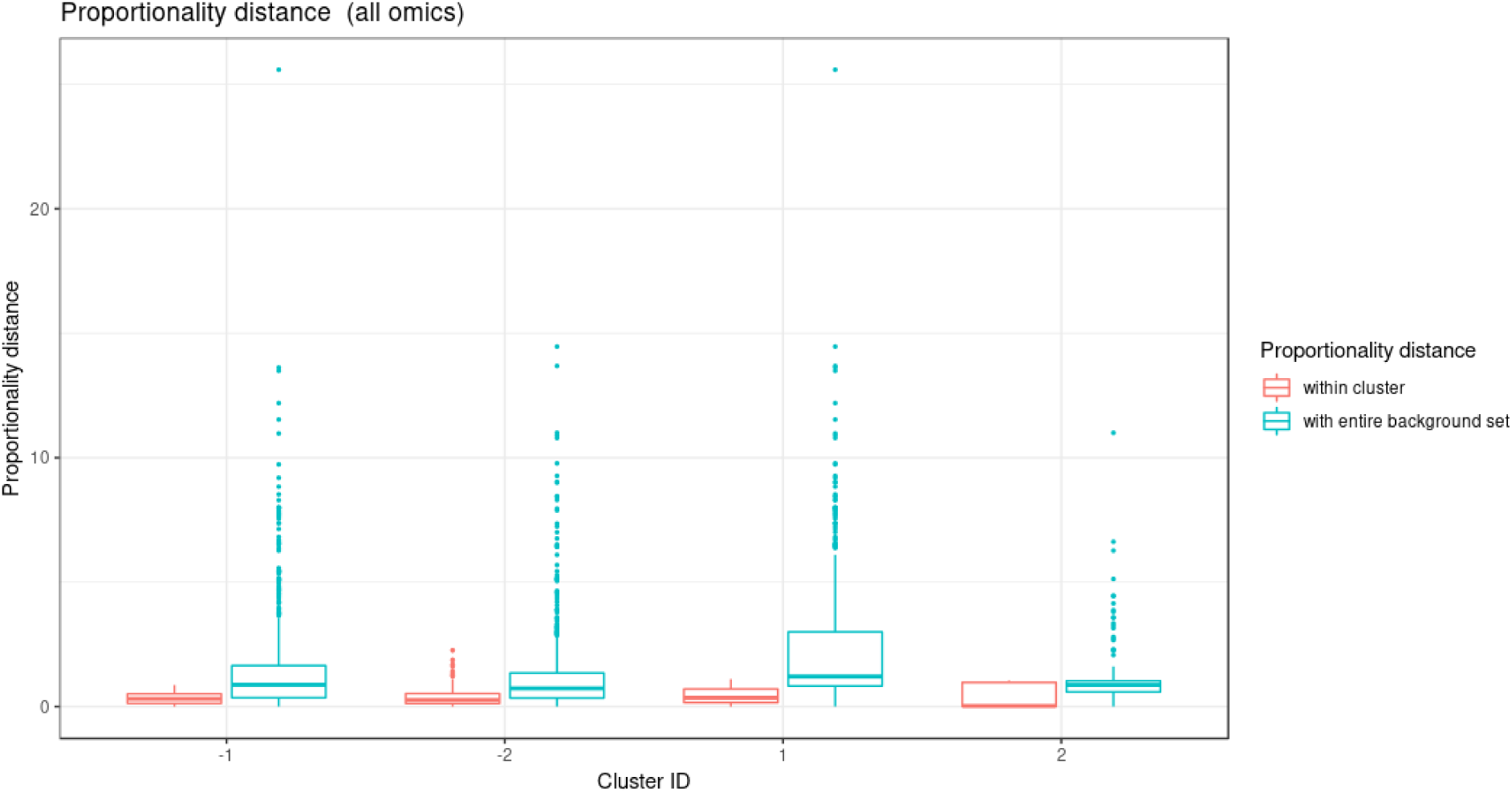
Waste degradation study: Proportionality distance per cluster identified with multiblock sparse PLS. The distance was calculated between each pair of profiles within a given cluster and with the entire background set (outside a given cluster), for all omics.

Two performance variables (methane and carbon dioxide production) were assigned to cluster 1 (component 1 negative). This result is biologically relevant, as biogas is the final output of the AD reaction and is known to be associated with microbial activity and growth. Moreover, it is produced by archaea, such as *Methanosarcina*, also selected in this cluster. The proportionality distance between this OTU and methane was very low (*φ*_*s*_ = 0.25, Supplementary Material S5) confirming a strong association. Cluster 1 therefore represented the progress of the degradation process. In Cluster 2 (component 1 positive), we identified acetate produced by bacteria in the early days of the incubation and consumed by archaea (Cluster 1) to produce biogas. It was logically negatively associated to cluster 1 representing the progress of the degradation. Propionate was assigned to cluster 3 (component 2 positive). Its degradation was delayed compared to the molecule of cluster 1. It was expected as, for thermodynamical reasons, its degradation usually only starts when all acetate is degraded (Chapleur *et al.*, 2014). It was biologically relevant to find it associated with Hydrocinnamic and 3,4-Dihydroxyhydrocinnamic acids, which are also difficult to degrade. Cluster 4 (component 2 negative) was composed of only OTUs and metabolites and was similar to the one obtained with sPLS.

In summary, our framework allowed us to integrate different omic datasets measured longitudinally, and identify subsets of relevant microorganisms that were highly associated with metabolites abundance and performance measures through the biodegradation process. These analyses constitute a first step towards generating novel hypotheses about the biological mechanisms underpinning the dynamics in anaerobic digestion.

## 5 Discussion

Advances in technology and reduced sequencing costs have resulted in the emergence of new and more complex experimental designs that combine multiple omic datasets and several sampling times from the same biological material. Thus, the challenge is to integrate longitudinal, multi-omic data to capture the complex interactions between these omic layers and obtain a holistic view of biological systems. In order to integrate longitudinal data from microbial communities with other omics, meta-omics or other clinical variables, we proposed a data-driven analytical framework to identify highly associated temporal profiles between these multiple and heterogeneous datasets.

The application of this method allows the identification of similar expression profiles within a particular dataset (e.g. infant gut microbiota development study) but also across heterogeneous data types (16S amplicon microbiome data, metabolomics, chemical data in the waste degradation study). The clustering of longitudinal profiles helps identify groups of biological entities that may be functionally related, and thus generate novel hypotheses about the regulatory mechanisms that take place within the ecosystem.

In the proposed framework, the microbial counts of the microbiota’s constituent species are normalized for uneven sequencing library sizes and compositional data. Modelling with linear mixed model splines enables us to reduce the dimension of the data across the different biological replicates and take into account the individual variability due to either technical or biological sources. This approach also enables us to compare data analyzed at different time points (e.g. the waste degradation study). Lastly, we clustered the data using multivariate dimension reduction techniques on the spline models that further allowed integration between different data types, and the identification of the main patterns of longitudinal variation.

Ribicic *et al.* (2018) proposed an approach similar to ours, but they applied individual PCA or sPCA on each dataset (chemical loss and microbial community) after local polynomial regression modelling. Integration was performed in a second stage of the analysis with PLS by using hierarchical clustering (Cluster Image Maps visualization) to identify correlations between the two datasets. In comparison, we offer a more complete framework that accommodates complex scenarios, across several omics, across replicates, and handles compositional data. The LMMS allows for the modelling of expression over time for each compound across biological replicates while taking into account the overall individual variability. We use sPCA, sPLS and block sPLS as clustering means by leveraging on the loading vectors from these methods while selecting meaningful profile signatures.

Integrating different types of microbiome longitudinal data (e.g. abundance, activity, metabolic pathways, or macroscopic output) can be naively performed by concatenating all datasets. However, we showed that this approach was unsuccessful at selecting a sufficiently large number of profiles of different types, and thus did not shed light on the holistic view of the ecosystem dynamics (bioreactor study). Our integrative multivariate methods sPLS and block sPLS were better suited for the integration task, as they do not merge but rather statistically correlate components built on each dataset, and thus avoid unbalance in the signature when one dataset is either more informative, less noisy, or larger than the other datasets.

When compared with fPCA, that uses either *k*-CFC or EM clustering algorithms, we showed that our approach led to better clustering performance. In addition, the sparse multivariate approaches sPCA and block sPLS enabled the identification of key profiles to improve biological interpretation. Note however that fPCA might be better suited than our approach for a large number of time points, as we discuss next.

We have identified several limitations in our proposed framework. First, a high individual variability between biological replicates limits the LMMS modelling step, resulting in simple linear regression models to fit the data. Whilst a straight line model may accurately describe temporal dynamics, it could also be due to a poor quality of fit. We have implemented the Breusch-Pagan test to address this issue. Alternatively, in the case of a very high inter-individual variability that prevents appropriate smoothing, one could consider *N of One* analyses as proposed by Gerber *et al.* (2012); Äijö *et al.* (2017) with time dynamical probabilistic models.

Second, a large number of time points can result in the modelling of noisy profiles and clusters, often due to high individual variability. Highly variable and vastly different profiles can also be difficult to cluster appropriately. Therefore, this framework is recommended when the number of time points remains small (5-10) and when regular and similar trends are expected from the data.

Third, even though our simulation results showed that the LMMS interpolation of missing time points did not seem to impact clustering, the overall performance of the approach would be optimal for regularly spaced time points in the omics longitudinal experiments.

Fourth, we have not fully addressed the issue of analyzing time-course compositional data. Indeed, when working with relative abundances (TSS), fluctuations in the abundance of a particular microorganism might result in spurious fluctuations in the abundance of other microorganisms. This issue is not specific to microbiome data only, as other sequenced-based data are intrinsically compositional (Gloor *et al.*, 2017). Thus, when looking for associations between longitudinal profiles, the optimal solution could be to analyze absolute abundances. However, such data require spike-ins and are currently rarely available. Badri *et al.* (2018) have investigated normalization strategies and their effect in correlation analysis but for a single time point, while Metwally *et al.* (2018) proposed three normalization strategies that ignore the compositionality data problem. No method for longitudinal compositional data analysis has been proposed as yet. The proportionality measure proposed by Lovell *et al.* 2015 is a promising solution to reduce spurious correlations. However, it has not been developed for longitudinal problems, and the metric is not suitable in our context to perform variable selection. Instead, we chose to use the proportionality distance as a post-hoc evaluation in our framework, not only to reduce potential spurious associations between profiles assigned in each cluster, but also to improve and help interpretation with respect to proportional and relative abundance of the profiles.

Finally, our framework does not include time delay analysis, even though dynamic delays between different types of molecules (e.g. DNA, RNA, or metabolites) can be expected. For example, 16S data describes the abundance of the microorganisms, with metabolites as the consequence of their activity, and performance as the macroscopic resulting output. Potential delays between these molecules can be detected using other techniques, such as the Fast Fourier Transform approach from Straube *et al.* (2017), and will be further investigated in our future work.

To summarize, we have proposed one of the first computational frameworks to integrate longitudinal microbiome data with other omics data or other variables generated on the same biological samples or material. The identification of highly associated key omics features can help generate novel hypotheses to better understand the dynamics of biological and biosystem interactions. Thus, our data-driven approach will open new avenues for the exploration and analyses of multi-omics studies.

## 6 Additional Requirements

### Author Contributions

All authors contributed to the design of the study; AB and OC performed the statistical analyses; AB, OC and KALC wrote the manuscript. All authors read and approved the submitted version.

### Funding

KALC was supported in part by the National Health and Medical Research Council (NHMRC) Career Development fellowship (GNT1159458). KALC and OC scientific travels were supported in part by the France-Australia Science Innovation Collaboration (FASIC) Program Early Career Fellowships from the Australian Academy of Science. AD was supported by Research and Innovation chair L’Oreal in Digital Biology.

## Acknowledgments

We thank Angéline Guenne for analytical support with GC-MS analysis, Kodjovi Dodji Mlaga for the biological interpretations of the infant study, and Zoe Welham for proof-reading the manuscript. We thank the reviewers for their constructive comments.

## Supplemental Data

Available as online document.

## Data Availability Statement

Infant gut microbiota phylochip raw data can be found in Palmer *et al.* (2007). The microbiome and performance datasets for the bioreactor study can be found in Poirier and Chapleur (2018), metabolomic data is available on request. In-house scripts and code to conduct both case study analysis, are available in a Github public repository: https://github.com/abodein/timeOmics

## A Linear Mixed Model Splines (LMMS) models

The first model assumes the response is a straight line not affected by individual variation. Let *y*_*ij*_(*t*_*ij*_) be the taxa normalized count for individual (or biological replicate) *i* at time *t*_*ij*_, where *i* = 1, 2, *…, n, j* = 1, 2, *…, m*_*i*_, *N* is the sample size and *m*_*i*_ is the number of observations for individual *i* for the given taxa. A simple linear regression of abundance *y*_*ij*_(*t*_*ij*_) on time *t*_*ij*_, with the intercept *β*_0_ and slope *β*_1_ is estimated via ordinary least squares:

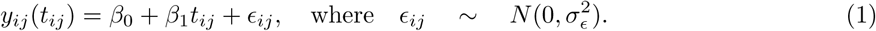

As nonlinear response patterns are commonly encountered, a second model uses a spline truncated line basis as proposed by Durban Durbán *et al.* (2005) to model a curve:

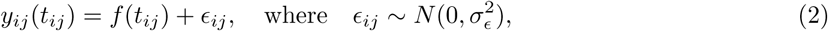

where *f* represents a penalized spline which depends on a set of knot positions *κ*_1_, *…, κ*_*K*_ in the range of {*t*_*ij*_}, some unknown coefficients *u*_*k*_, an intercept *β*_0_ and a slope *β*_1_, i.e.

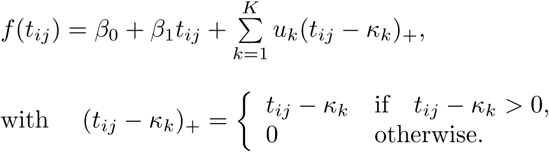

The choice of the number of knots *K* and their positions influences the flexibility of the curve. As proposed by Ruppert (2002), we estimate the number of knots based on the number of measured time points *T* as 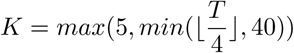, placing the knots *κ*_1_*…κ*_*K*_ at quantiles of the time interval of interest.

A third model accounts for individual variation in Eq. (3) with the addition of a subject-specific random effect *U*_*i*_ to the mean response *f* (*t*_*ij*_). We assume *f* (*t*_*ij*_) to be a fixed (yet unknown) population curve, *U*_*i*_ is treated as a random realisation from an underlying Gaussian distribution independent from the previously defined random error *ϵ*_*ij*_. The individual curves are expected to be parallel to the mean curve as we assume the subject-specific random effects to be constant over time:

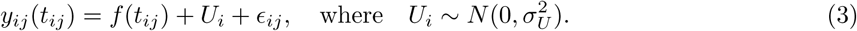

The final and fourth model is an extension to Eq. (3) that assumes individual deviations are straight lines, where individual-specific random intercepts *a*_*i*0_ and slopes *a*_*i*1_ are fitted:

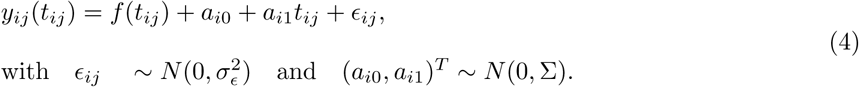

Here we assume independence between the random intercept and slope, so the covariance matrix for the random effects Σ is diagonal.

## B Supplementary Figures and Tables

**Figure S1:**
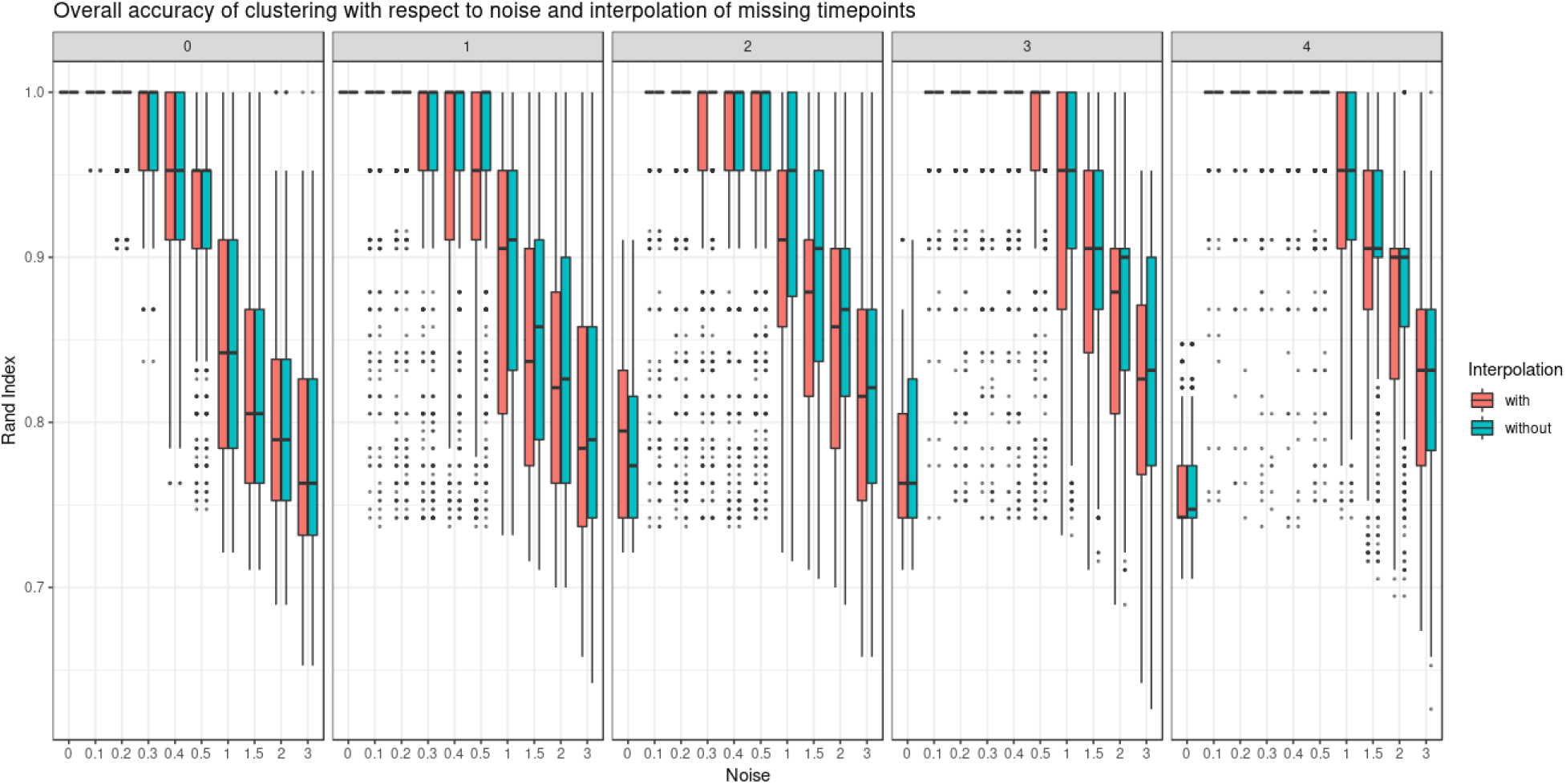
Simulated study: Overall compactness of assigned clusters when time points are missing with the Rand index.

**Figure S2:**
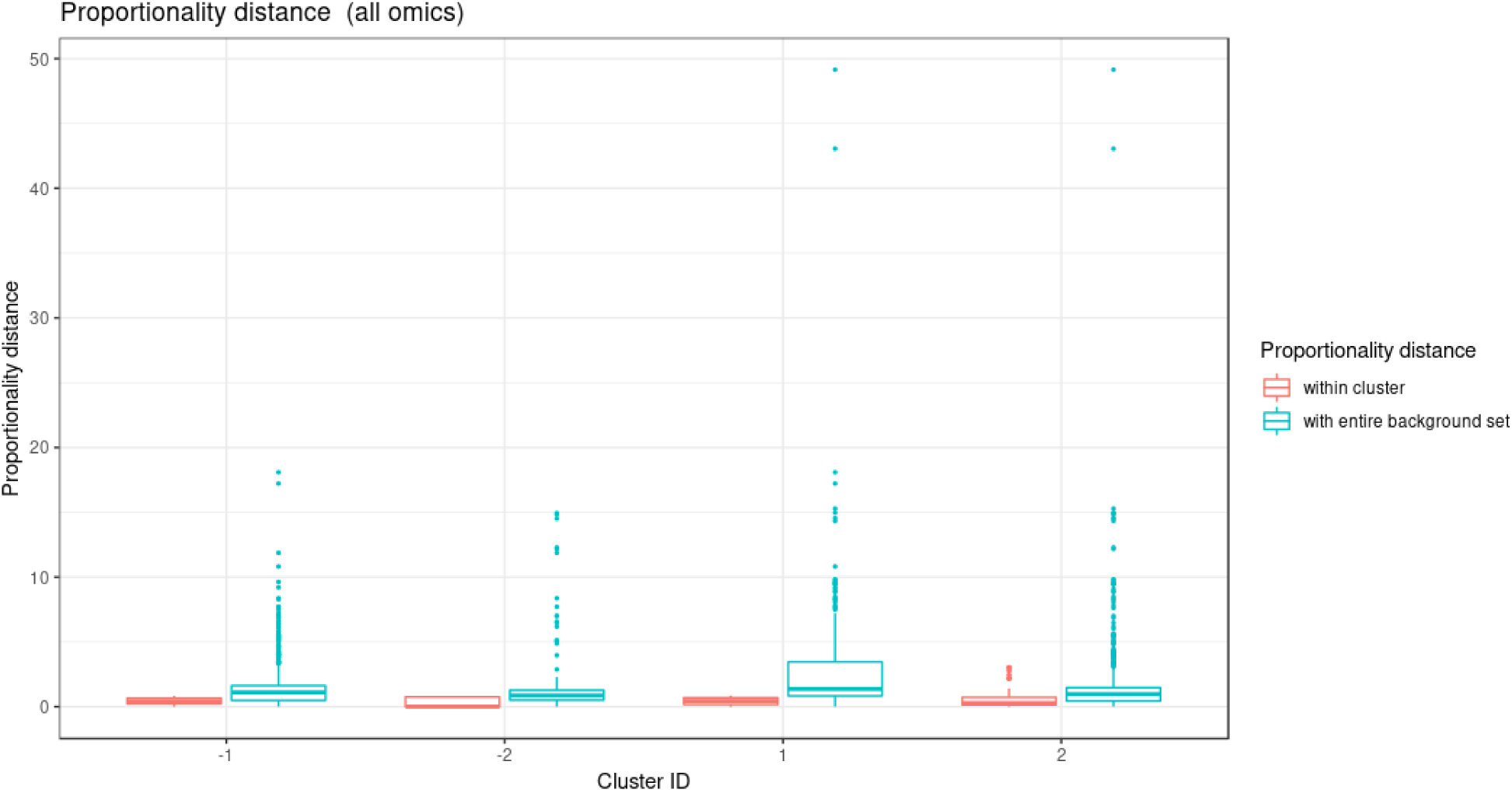
Waste degradation study: Proportionality distance per cluster identified with sparse PLS. The distance was calculated between each pair of profiles within a given cluster and with the entire background set (outside a given cluster).

**Figure S3:**
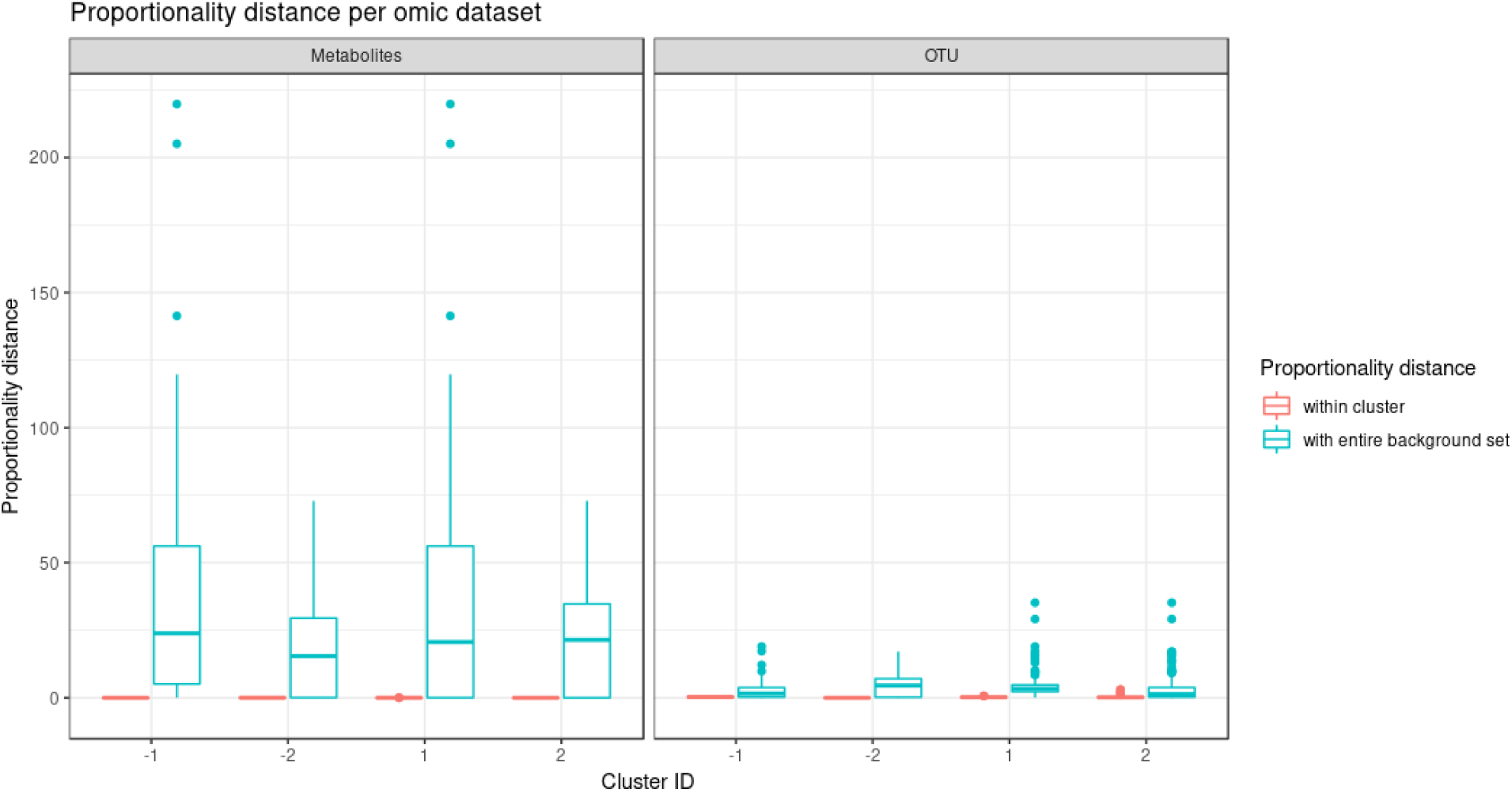
Waste degradation study: Proportionality distance per cluster and per omic dataset on the profiles identified with sparse PLS clustering. The distance was calculated between each pair of profiles of the same type of omics data type within a given cluster and with the entire background set (outside a given cluster). Distances are displayed per omic dataset.

**Figure S4:**
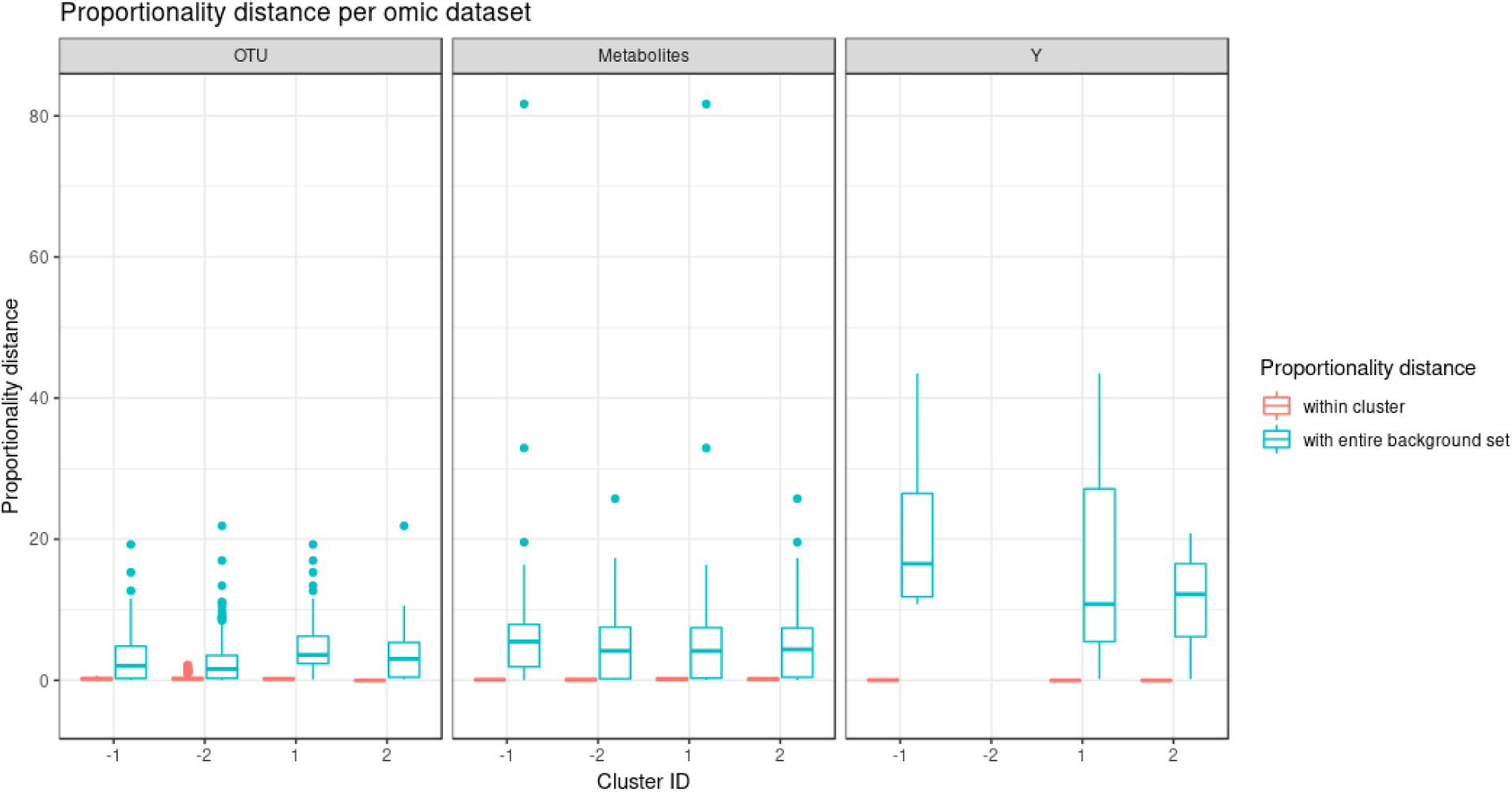
Waste degradation study: Proportionality distance per cluster and per omic dataset identified with multiblock sparse PLS clustering. The distance was calculated between each pair of profiles of the same type of omics data type within a given cluster and with the entire background set (outside a given cluster). Distances are displayed per omic dataset.

**Figure S5:**
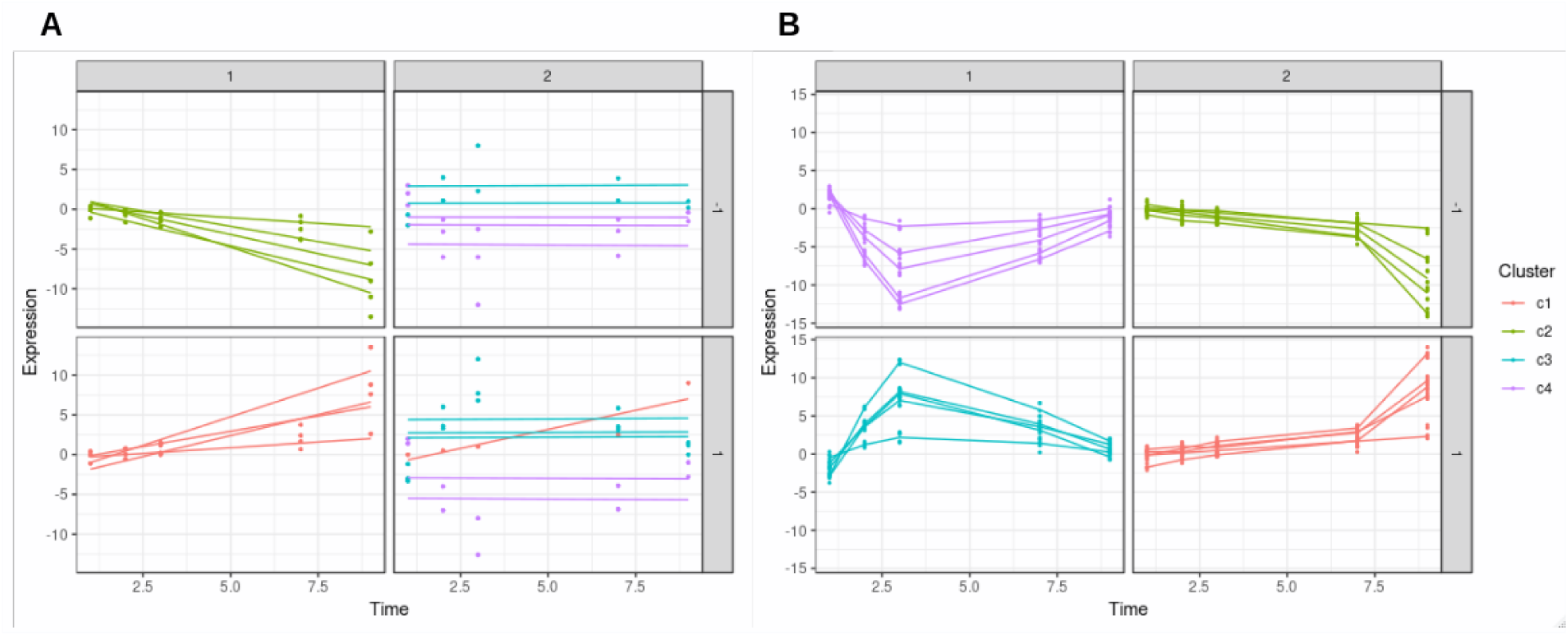
Simulation study: Clustering of simulated profiles with 4 missing time points, when noise = 0. Colors represent the ground truth cluster. (**A**) When no time points are missing, LMMS mostly modelled straight lines, resulting in a poor clustering assignation. (**B**) When time points are missing, LMMS modelled splines resulted in better clustering.

**Figure S6:**
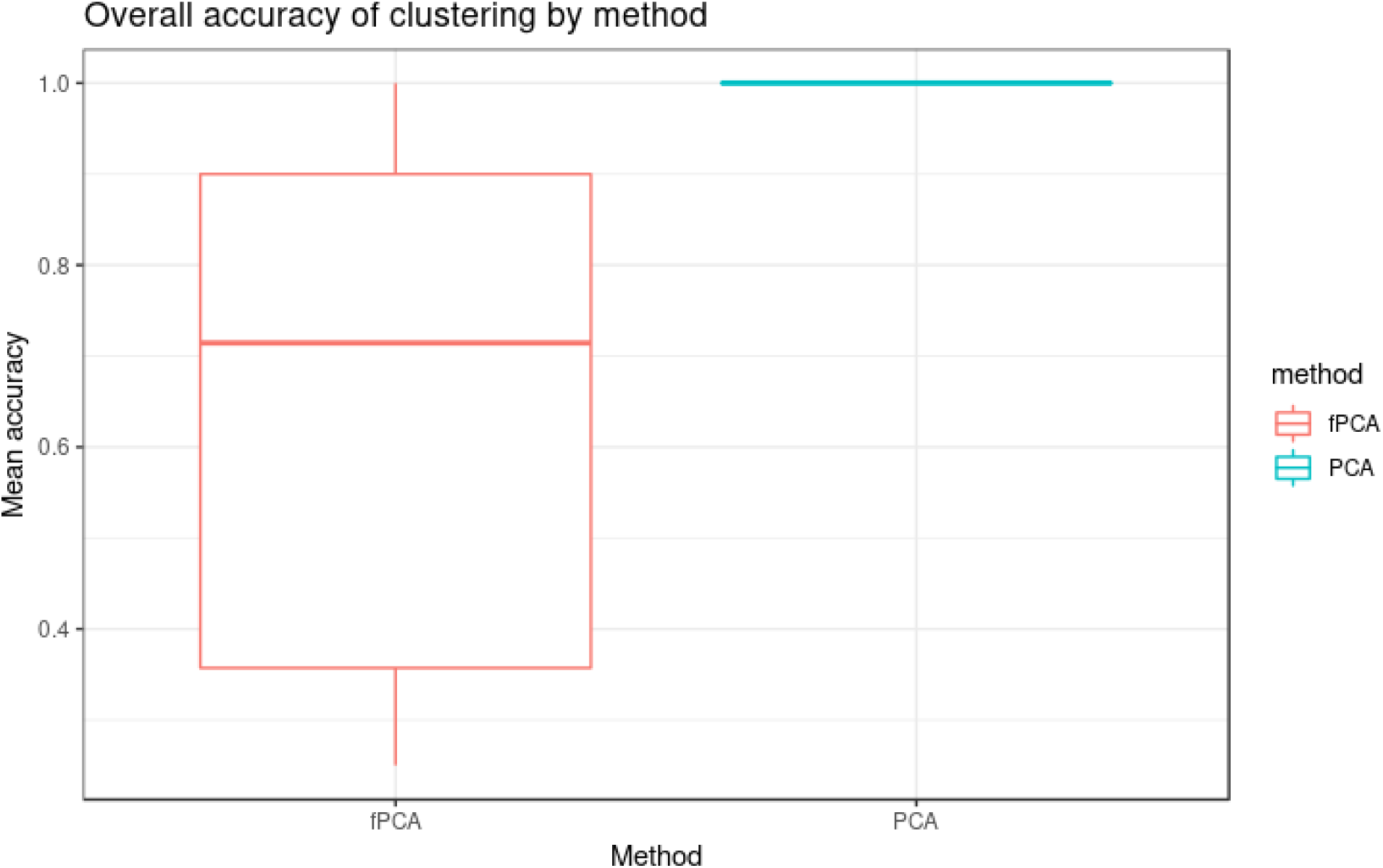
Simulation study: Overall accuracy of PCA and fPCA clustering with no noise. We compared the ability of PCA and fPCA to correctly assign the simulated profiles in their respective reference clusters. Without noise, fPCA clustering led to a poorer accuracy compared to PCA.

**Table S1:** Infant gut microbiota development for vaginal delivery. Proportionality distance for clusters identified with PCA. The median distance between all pairs of profiles, within cluster and with the entire background set (outside a given cluster) is reported. A Wilcoxon test p-value assesses the difference between the medians.

| cluster | median within cluster | median outside cluster | Wilcoxon test P-value |
| --- | --- | --- | --- |
| 1 (comp 1 positive) | 0.64 | 1.3 | $2.99 * 10^{-149}$ |
| -1 (comp 1 negative) | 0.2 | 1.44 | $\leq 10^{-149}$ |
| 2 (comp 2 positive) | 0.44 | 1.11 | $2.59 * 10^{-15}$ |
| -2 (comp 1 negative) | 0.35 | 0.95 | $2.75 * 10^{-19}$ |

**Table S2:** Infant gut microbiota development for C-section delivery. Proportionality distance for clusters identified with PCA. The median distance between all pairs of profiles, within cluster and with the entire background set (outside a given cluster) is reported. A Wilcoxon test p-value assesses the difference between the medians.

| cluster | median within cluster | median outside cluster | Wilcoxon test P-value |
| --- | --- | --- | --- |
| 1 (comp 1 positive) | 0.11 | 1.36 | $3.46 * 10^{-80}$ |
| -1 (comp 1 negative) | 0.29 | 0.96 | $1.36 * 10^{-15}$ |
| 2 (comp 2 positive) | 0.67 | 1.5 | $1.02 * 10^{-103}$ |
| -2 (comp 2 negative) | 0.16 | 1.28 | $1.21 * 10^{-95}$ |

**Table S3:** Waste degradation study: Proportionality distance for clusters identified with multiblock sparse PLS. The median distance between all pairs of profiles, within cluster and with the entire background set (outside a given cluster) is reported. A Wilcoxon test p-value assesses the difference between the medians.

| cluster | median within cluster | median outside cluster | Wilcoxon test P-value |
| --- | --- | --- | --- |
| 1 (comp 1 positive) | 0.36 | 1.21 | $1.25 * 10^{-55}$ |
| -1 (comp 1 negative) | 0.31 | 0.88 | $9.84 * 10^{-36}$ |
| 2 (comp 2 positive) | 0.03 | 0.87 | $3.34 * 10^{-5}$ |
| -2 (comp 2 negative) | 0.27 | 0.73 | $7.18 * 10^{-22}$ |

Table S4:Infant gut microbiota development study: Proportionality distance between all pairs of OTUs. Clustering assignments from PCA and sPCA are indicated, along with whether the OTU was selected with sPCA. First sheet: vaginal data, Second sheet: for C-section data. External file ‘propr_*g*_*ut.xls*′

Table S5:Waste degradation study: Proportionality distance between all pairs of entities (OTUs, metabolites and performance data). Clustering assignments from either PLS (first sheet, integration of OTUs and metabolites) or multiblock PLS (second sheet, integration of all three datasets). Selection with sPLS or multiblock sPLS are indicated. External file ‘propr_*b*_*iowaste.xls*′.

